# Orthogonal Modes of Gene Expression Evolution Shape Human Neocortical Development and Disease Vulnerability

**DOI:** 10.64898/2025.12.11.693619

**Authors:** Xuanhao D. Zhou, Yuki Yamauchi, Kazuo Emoto, Ikuo K. Suzuki

## Abstract

The neocortex is responsible for higher-order cognitive abilities such as language, abstract reasoning, and executive function—capacities that are particularly advanced in humans. To elucidate the molecular foundations of neocortical evolution in the human lineage, it is essential to examine how conserved gene repertoires have undergone expression changes relative to other mammals. The expression changes in conserved genes are widely regarded as key drivers of the phenotypic evolution of human-specific traits. In this study, we performed a comprehensive comparative single-cell transcriptomic analysis of fetal neocortical development across four mammalian species: human, macaque, mouse, and ferret. To validate human-specific expression changes, we further analyzed brain organoids derived from both human and chimpanzee stem cells. Genes exhibiting human-specific expression shifts were systematically classified along three orthogonal dimensions: overall expression level (Human Level Distinctive; HLD), temporal expression trend (Human Trend Distinctive; HTD), and differentiation lineage specificity (Human Differentiation trajectory Distinctive; HDD). HLD genes were frequently enriched for long introns and located near Human Accelerated Regions (HARs), and showed pronounced upregulation in humans. These genes were strongly associated with neurodevelopmental disorders such as autism spectrum disorder and developmental delay, as well as with megalencephaly and glioblastoma. HTD genes, in contrast, exhibited a unique pattern in humans, peaking early in development and subsequently declining—opposite to the steadily increasing trends observed in other species. These genes were significantly enriched for oxidative phosphorylation and ribosomal functions, pointing to a temporally restricted elevation in biosynthetic activity in early human corticogenesis. HDD genes displayed a marked shift in lineage-specific expression: cilia-related genes that are typically expressed in apical progenitors in non-human species were instead highly expressed in outer radial glia (oRGs) in humans. This spatial reorganization of ciliary gene activity suggests an oRG-specific adaptation in signaling architecture. Together, these results highlight the diversity of regulatory changes that have shaped human cortical development. Distinct classes of gene expression evolution—mediated in part by HARs—appear to have contributed not only to the expansion and increased complexity of the human neocortex, but also to its heightened vulnerability to neurodevelopmental and oncogenic pathologies. The identified human distinctive genes will be the target of future experimental verification to elucidate the precise molecular mechanisms regulating human-specific aspects of cortical development.

## Introduction

The neocortex is the locus of higher-order cognitive abilities such as language, abstract reasoning, and executive function, which are especially elaborated in humans. It dominates human brain volume and is composed of a vast diversity of neurons and glial cells organized into highly complex circuits. While the fundamental cytoarchitecture and cellular diversity of the neocortex are broadly conserved among mammals, substantial differences in size and cellular abundance underlie species-specific cognitive capacities^1–7^ (Fig.1A). Since the vast majority of neocortical cells are generated during embryonic development, interspecies differences in cortical structure are ultimately rooted in evolutionary changes in developmental programs.

During embryogenesis, the neocortex arises from the dorsal telencephalon, specified once brain regionalization along the embryonic axes is complete. Within this territory, neuroepithelial stem cells initially expand the progenitor pool through symmetric divisions. These cells then transition into ventricular radial glia (vRGs), which begin producing excitatory neurons (ExN). Nascent neurons migrate radially toward the pial surface to form the cortical plate. vRG progenitors contribute to neurogenesis both directly and indirectly, via intermediate progenitors such as intermediate progenitors (ExN-IPC) and outer radial glia (oRGs, also known as basal radial glia), both residing in the subventricular zone (SVZ). These intermediate progenitors amplify neuronal output through additional rounds of proliferation. As neurogenesis terminates, gliogenesis initiates with the generation of astrocytes and oligodendrocytes, which arise through lineage transitions involving tRGs, oRGs, and dedicated glial intermediate progenitors (glia-IPCs). These immature glial cells, immature astrocytes (Ast-imma) and oligodendrocyte precursor (OPC), eventually exit the cell cycle and undergo terminal differentiation. Thus, neuro- and gliogenesis constitute a continuous cascade of cell-state transitions originating from vRG population (Fig.1B).

Although the basic sequence of cell fate transitions is highly conserved across mammals, differences in the timing, amplification, and spatial dynamics of progenitor populations have contributed to the diversification of cortical architectures^1–3,5,8^. Gyrencephalic species such as primates and carnivores exhibit enhanced proliferation of neuroepithelial stem cells and an expanded surface area, in contrast to lissencephalic rodents with smaller, smoother cortices. Another hallmark of cortical expansion in gyrencephalic species is the formation of distinct tRG and oRG populations, which emerge during mid-neurogenesis when the radial scaffold is physically severed in the SVZ^9,10^. This leads to the spatial segregation of progenitor niches—ventricular-facing tRGs and basally located oRGs, whereas rodents maintain a unified vRG population throughout development with the presence of a minor oRG population. oRG progenitors produce hundreds of neurons to locally expand neuronal output in the future gyrus in the large-brained gyrencephalic animals^11–13^. In addition, primates, especially humans, exhibit significantly prolonged neurogenic and gliogenic time windows compared to rodents; despite a relatively minor difference in cell-cycle durations, the extended developmental timeframe enables the generation of substantially more neurons and glia in species (Supplementary Fig.1A)^1,5,8^.

At the molecular level, the genetic programs governing cortical development are deeply conserved, enabling cross-species comparisons of homologous cell types using shared marker genes and signaling pathways. However, a growing number of studies have uncovered human-specific genetic mechanisms that modulate cortical development^1,14–19^. Several groups, including our own, have identified human-specific genes that emerged in the human lineage and contribute to various aspects of brain development^20–28^. While such gene births are relatively rare, changes in conserved gene repertoire are believed to be more common. Examples include human-specific coding mutations in genes regulating cortical development^29–31^, such as FOXP2^32^. In line with the seminal hypothesis^33^, which posited that regulatory changes in conserved genes are central to human-specific traits, a number of transcriptomic and epigenomic studies have revealed extensive changes in gene expression and regulation during human cortical development^34–37^. Recent analyses of epigenetic modifications have further identified human-accelerated regions (HARs) with enhancer activity, including functional validation in the case of HARE5, which upregulates FZD8 expression and expands cortical size^38–42^. Collectively, these studies underscore how evolutionary alterations in gene regulatory landscapes can reshape developmental trajectories and, ultimately, neural architecture.

Importantly, evolutionary changes in gene expression can manifest in multiple distinct forms^43^. Some genes undergo “heterometric” shifts, characterized by changes in overall expression magnitude. Others display “heterochronic” differences, wherein the temporal dynamics of gene expression are modified. Still others exhibit “heterotopic” changes, in which genes are newly expressed in different developmental lineages or trajectories. These modes of regulatory evolution are not mutually exclusive; a single gene theoretically exhibit more than one type of change. Each category likely carries unique developmental and evolutionary consequences, yet the relative contribution of each to human brain evolution remains largely unexplored. To our knowledge, no prior study has systematically classified these distinct forms of gene expression changes during fetal neocortical development in a multi-species comparative context.

Advances in single-cell transcriptomics now permit single cell resolution comparisons of gene expression across species and developmental timepoints, enabling accurate identification of homologous cell types and their differentiation trajectories. However, because developmental cell states represent a continuum, relying solely on discrete marker-based annotations may obscure subtle but important interspecies differences. Moreover, the majority of comparative studies have focused on pairwise comparisons—typically between human and one other species—limiting the ability to infer ancestral states and confidently identify human-specific changes.

To overcome the limitations of previous studies, we performed an integrated comparative single-cell transcriptomic analysis of neocortical development across four mammalian species—human, macaque, mouse, and ferret—with the aim of identifying human-specific gene expression changes. These findings were further validated through comparisons between human and chimpanzee using *in vitro* cortical organoid models (Fig. 1A). Although cortical organoids are not perfect surrogates for the fetal brain—owing to their lack of vasculature and non-neural tissues—they remain valuable systems for evolutionary studies, as they faithfully recapitulate the full spectrum of vRG-derived lineages present during corticogenesis in both humans and great apes^44,45^, whose embryonic tissues are practically inaccessible. Focusing on both neurogenic and gliogenic phases of development, we systematically quantified interspecies divergence in gene expression along three orthogonal axes: overall expression levels, temporal expression dynamics, and differentiation trajectory specificity. By comparing humans to three distinct non-human species, we robustly defined human-distinctive gene sets for each axis of regulatory divergence, providing a comprehensive framework to dissect the molecular underpinnings of human brain evolution.

**Figure 1.**
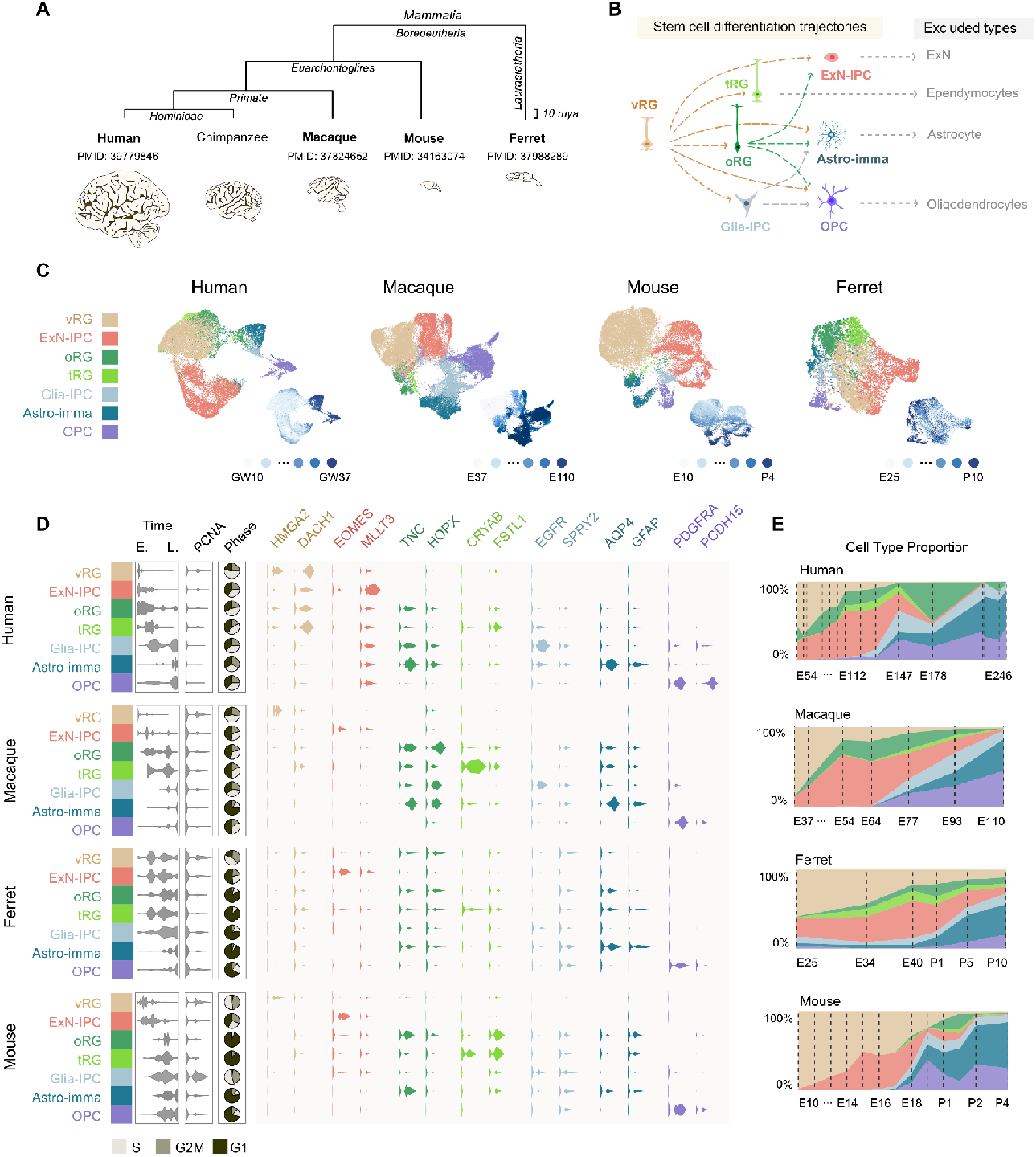
Integrated Progenitor Dynamics from Four Mammalian Species Neocortical Development. **(A-B)** Schematic of the study design, integrating single-cell RNA sequencing (scRNA-seq) data from fetal neocortical tissues of human, macaque, mouse, and ferret, spanning neurogenesis to early gliogenesis. Bulk RNA-seq data from human and chimpanzee cerebral organoids are included for validation of human-specific expression trends. **(C)** UMAP embedding of single-cell datasets, with cells classified into seven distinct cell types for each species. Developmental stage progression is visualized by a color gradient from white (early) to dark blue (late). **(D)** Temporal dynamics, proliferative capacity (PCNA expression), predicted cell cycle phase, and canonical marker gene expression for each cell type across species. **(E)** Proportional changes in cell type composition across developmental stages. Dashed lines indicate stages sampled for single-cell RNA sequencing.

## Results

### Integrated transcriptomic analysis of fetal neocortex across multiple mammalian species

To investigate the evolutionary divergence of gene expression during neocortical development, we obtained publicly available single-cell transcriptomics datasets of fetal neocortical tissues from four mammalian species: human^46^, macaque^47^, mouse^48^, and ferret^9^ and performed an integrated comparative analysis (Fig.1A). Samples were collected from matched developmental windows spanning the onset of neurogenesis through the early phase of gliogenesis (Supplementary Fig.1). In select analyses, we also incorporated bulk RNA sequencing data of human and chimpanzee cerebral organoids prepared in house to validate human-specific expression trends observed *in vivo*.

To ensure robust dataset integration, we tested five methods (CCA, RPCA, Harmony, fastMNN, and SCT regression) and selected the optimal approach for within-species alignment based on biological and technical benchmarks (Supplementary Fig.2-Supplementary Fig.9). Cell-type annotations were based on the expression of conserved marker genes, enabling the identification of homologous cell populations despite differences in evolutionary lineage and cortical architecture (Fig.1A-B, marker conservation in Supplementary Fig.10 and metacell correspondence in Supplementary Fig.11). Given our interest in comparing dynamic expression change during progenitor expansion and continuous fate transitions, we focused exclusively on proliferative populations, vRG, ExN-IPC, oRG, tRG, Glia-IPC, Ast-imma, and OPC (Fig.1B). Thus, postmitotic neurons and fully differentiated glial cells were excluded from downstream analyses. We analyzed 30299, 51324, 25758 and 7698 proliferative cortical cells for human, macaque, mouse, and ferret in total.

In all four species, early developmental stages were dominated by vRGs (Fig.1C-E). As development progressed, the proportion of vRGs declined, giving way to increasing fractions of intermediate progenitors, including ExN-IPCs, oRGs, and tRGs (Fig.1E). At mid-developmental stages, glial progenitor populations such as glial-IPC, immature astrocytes (Astro-imma) and oligodendrocyte precursor cells (OPCs) began to emerge and increased steadily in abundance. While the temporal progression of cell-type composition followed a broadly conserved trajectory across all species, notable species-specific differences were observed. Most strikingly, mice exhibited a marked paucity of oRGs and tRGs compared to the other three species, in which these progenitor types were readily detectable. Additional interspecies differences were also evident, though some of these may reflect variation in tissue sampling strategies or developmental staging across species. Therefore, although global trends appear conserved, the biological significance of finer differences requires cautious interpretation.

### Identification of three classes of human distinctive gene expression changes

To systematically assess various modes of evolutionary divergence in gene expression patterns, we focused on a set of 10,683 orthologous genes that were commonly present across all four species. Our goal was to classify these genes based on the extent and nature of interspecies variation in their expression profiles during cortical development and to identify the genes that show distinctive expression signatures in humans. For each gene, we performed pairwise comparisons between species, examining every possible pair of the four animals. These comparisons were conducted along three orthogonal axes: (1) overall expression levels (Fig.2A), (2) temporal trend or dynamics of expression during development (Fig.2B), and (3) lineage-specific expression along differentiation trajectories (Fig.2C).

Differences in overall expression levels were quantified while correcting for potential biases due to uneven cell-type composition, which can otherwise distort interspecies comparisons. To address this, we performed logistic regression using cell type as a covariate, allowing us to control for compositional variability and isolate biologically meaningful differences. For each gene, we calculated the log fold change between species (Fig. 2A). For temporal dynamics, we normalized and centered gene expression values to focus specifically on developmental trajectories independent of expression magnitude. We then applied a Generalized Additive Model (GAM) along a pseudotime axis defined by supervised ordinal logistic regression with fixed developmental endpoints. Divergence between species was quantified as the Euclidean distance between their respective expression curves across pseudotime (Fig. 2B; see Methods for details). To quantify lineage-specific expression, we embedded cells into a lower-dimensional space based on transcriptional similarity, and constructed a transition matrix to estimate the cost of state transitions. Optimal differentiation trajectories were identified by minimizing this cost function. Jensen-Shannon Divergence (JSD) was then used to measure differences in the distribution of gene expression along these trajectories between species (Fig. 2C; see Methods for details).

**Figure 2.**
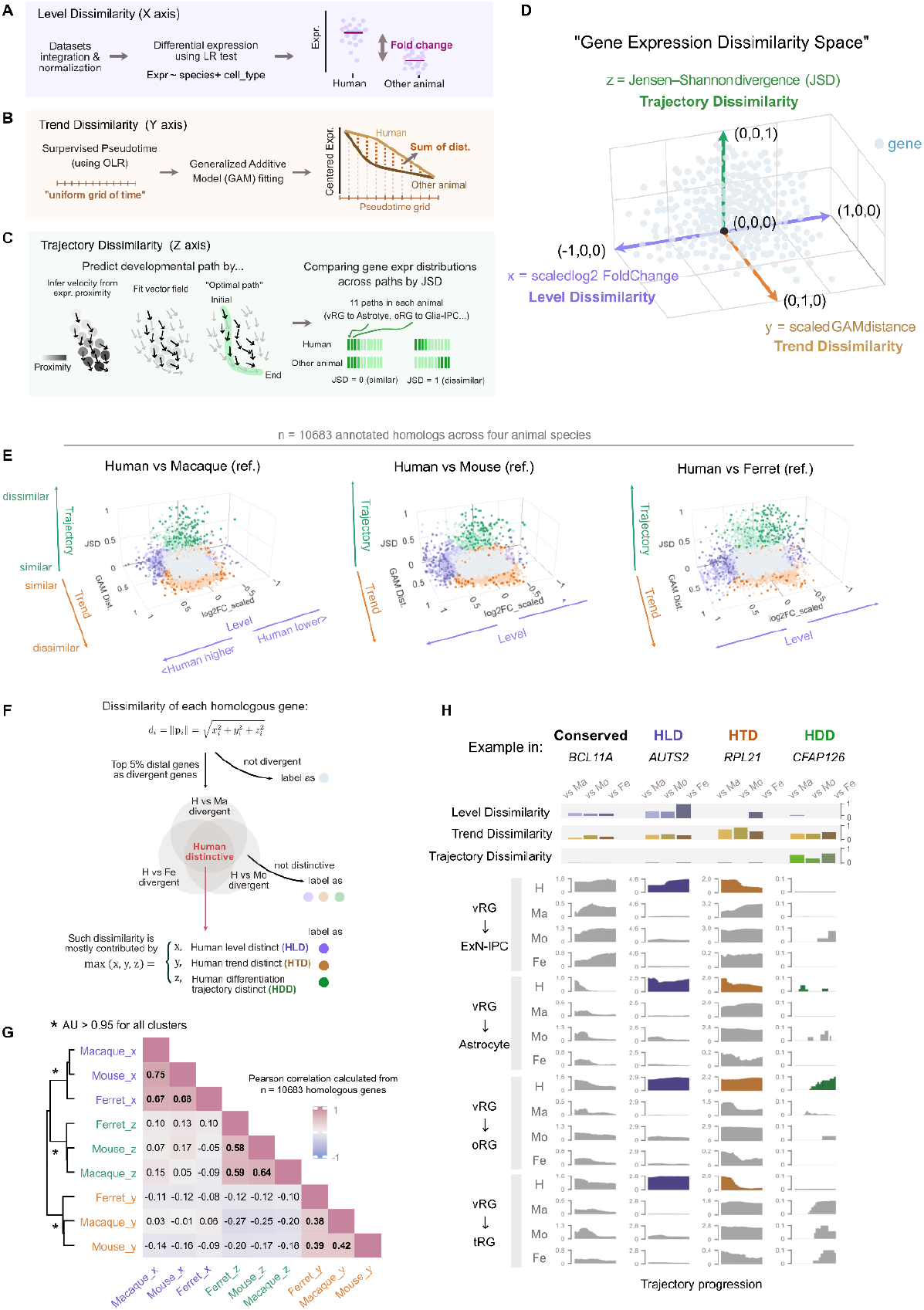
Mapping Cross-Species Divergence in Gene Expression Divergence Space. (A-D) Definition of Gene Expression Dissimilarity Space. A three-dimensional space with approximately orthogonal axes is defined as: Level (x), Trend (y), and Trajectory (z) dissimilarities. Each axis is scaled to the population distribution to ensure comparable metrics across dimensions. (B–D) Schematics illustrate the axis calculation methodology as well as computational approach for quantifying Level, Trend, and Trajectory dissimilarities. Refer to Methods for detailed procedures.(E–F) Definition of human divergent and distinctive genes. Definitions and annotations of human-specific divergent genes and their positions within the dissimilarity space.(G) Pearson correlation matrix of dissimilarity metrics across species comparisons. Asterisks (*) denote Approximately Unbiased (AU) p-values > 0.95, computed via pvclust.(H) Examples of genes from Human Level Divergence (HLD), Human Trend Divergence (HTD), and Human Trajectory Divergence (HDD), including their respective metrics and expression patterns along differentiation trajectories.

Crucially, both the temporal trend and lineage specificity metrics were constructed to be orthogonal to expression level, as they exclude magnitude information from their calculations. This enables the dissection of interspecies expression divergence across independent regulatory axes. Moreover, the lineage specificity score is independent of temporal trend because it is based on averaged expression values across all stages within each trajectory, and divergence is calculated from the distribution of these values across species using JSD.

To enable uniform comparisons, each axis was normalized relative to the distribution of all homologous genes. The expression level axis preserved directionality, scaling positive logFC from 0 to 1 and negative logFC from –1 to 0, maintaining biological interpretability. Temporal trend and trajectory specificity axes were scaled from 0 to 1, with 0 indicating conservation and 1 representing maximal divergence. These normalized scores positioned each gene within a three-dimensional coordinate system, termed the “Gene Expression Dissimilarity Space,” providing a rigorous, visualized framework to assess divergence in expression magnitude, developmental timing, and lineage specificity across species (Fig.2D). Visualization of this space clearly revealed how individual genes vary in expression magnitude (Level), developmental timing (Trend), and cell-lineage specificity (Trajectory) (Fig.2E). Collectively, these three axes accounted for approximately 90% of the total variation in cross-species expression profiles, evaluated by gradient Boosting machine learning (90.7% for human-macaque, 91.1% for human-ferret, and 89.4% for human-mouse comparisons, see methods for details). Expression level contributed most (46% macaque, 42% ferret, 46% mouse), followed by trajectories specificity (34%, 39%, 33%) and temporal dynamics (21%, 19%, 21%). We then applied this framework to specifically evaluate divergence between humans and the other three species. For each ortholog, we computed three species-difference scores based on the above metrics and performed correlation analysis (Fig.2G). Three key trends emerged: (1) the three metrics were largely independent, as scores on one axis showed minimal correlation with the others; (2) for each metric, results from human-vs-macaque, human-vs-mouse, and human-vs-ferret comparisons were highly concordant; and (3) the degree of correlation is highest within level distinctiveness, followed by trajectory and trend distinctivenesses, suggesting that expression magnitude is rather conservative among non-human mammals and only humans experienced significant changes during evolution. These results indicate that the three dimensions, expression level, temporal dynamics, and differentiation trajectory preference, represent orthogonal axes of evolutionary variation. Together, they capture the majority of interspecies expression divergence and enable precise categorization of genes with human-specific features along distinct regulatory dimensions.

To identify robust human-specific gene expression changes, we conducted systematic pairwise comparisons between humans and each of the three other species, macaque, mouse, and ferret, across the three dimensions defined previously (Fig.2E and F and Supplementary Fig.12). For each pairwise comparison, we extracted the top 5% of genes showing the most pronounced divergence in each axis (Fig.2F). Genes that were consistently identified as top 5% outliers in all three human–nonhuman comparisons were defined as human distinctive genes for that particular dimension, human level distinct (HLD) genes, human trend distinct (HTD) genes, and human differentiation trajectory distinct (HDD) genes, respectively (Fig.2H, Supplementary Table 1-Human distinctive genes).

### Human distinctive genes with altered expression levels

In the comparison of gene expression levels, we identified the top 5% most divergent genes (534 genes) in each of the three human–non-human species pairs: human–macaque, human–mouse, and human–ferret. Of these, 267 genes (45.5%) consistently exhibited human-specific expression shifts across all three comparisons and were classified as HLD genes. Gene ontology (GO) analysis identified that the terms “Glutamate receptor signaling pathway ( GO:0007215)”, “Cell-cell adhesion (GO:0098609)”, and “cAMP metabolic process (GO:0046058)” are found for these genes (Supplementary Fig.13). Weighted gene co-expression network analysis (WGCNA) further subdivided the 267 HLD genes into two primary clusters: a major group comprising genes that are upregulated in humans relative to non-human species, and a minor group of 25 genes that are downregulated in humans (Fig. 3A). The upregulated HLD genes were further resolved into three subgroups based on their expression profiles. The first group consists of genes with transient expression in progenitor populations, such as vRG, ExN-IPC, pRG, and tRG, and marked upregulation in glial lineages. GO analysis revealed a strong enrichment for the term “synaptic membrane” (GO:0097060) within this group. The second group is enriched for genes predominantly expressed in ExN-IPC and associated with “transmembrane receptor protein tyrosine phosphatase activity” (GO:0005001). The third group comprises genes specifically expressed in OPC and linked to the regulation of glutamatergic synaptic transmission (GO:0051966). WGCNA also identified transcription factor (TF) genes located at the central nodes of each network module, suggesting potential roles as upstream regulators. SOX5 and NR6A1 were identified as core TFs in the first group, while AFF3 and AUTS2 were implicated in regulating genes in the second group. For the human-downregulated gene module, ZBTB12 emerged as the central regulatory factor.

**Figure 3.**
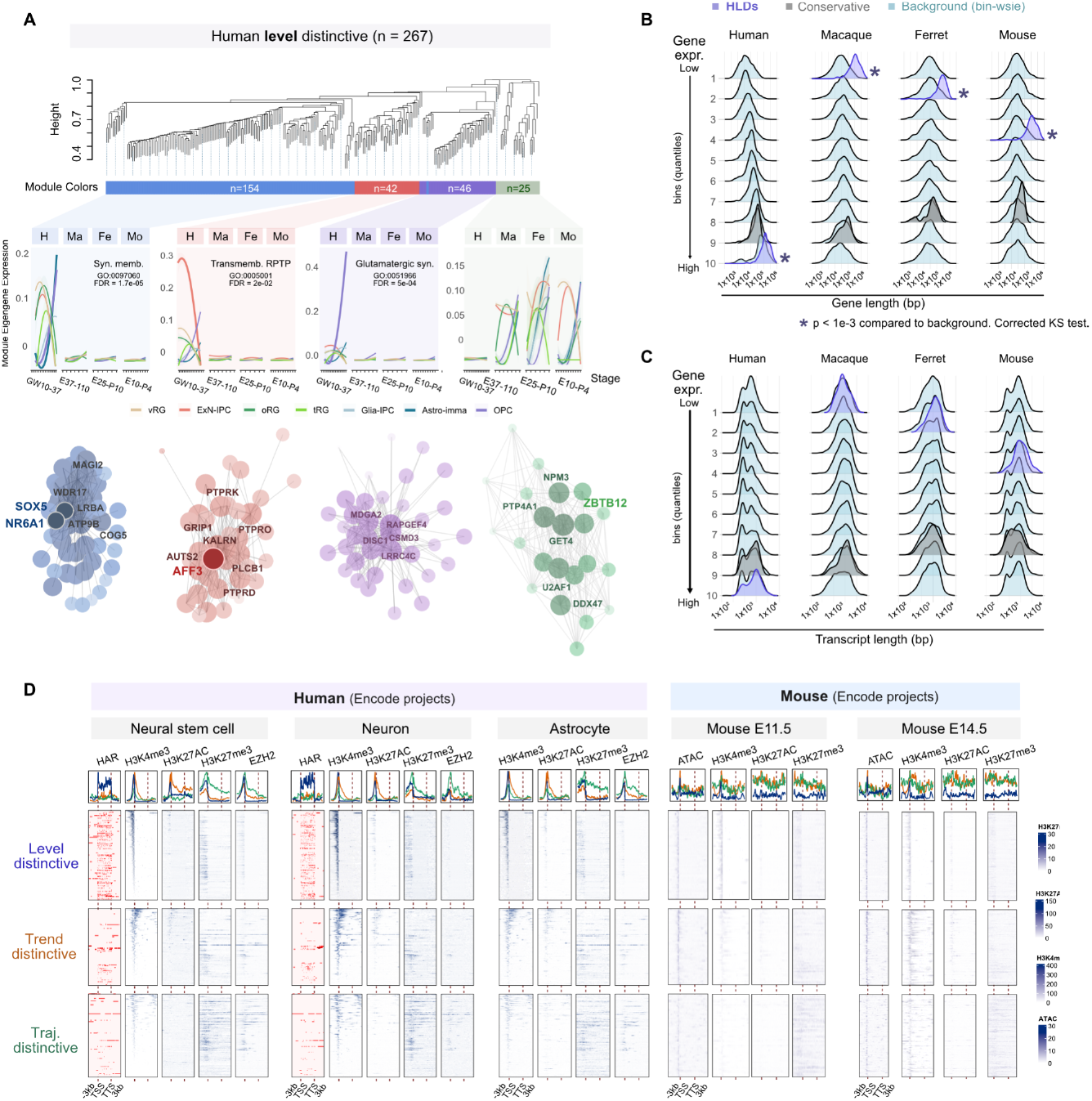
Elevated HLD Gene Expression Correlates with Extended Intron Length and Enriched HAR Interactions. **(A)** Weighted Gene Co-expression Network Analysis (WGCNA) of the HLD gene set identified four modules using default threshold settings (see Methods). Module eigengene expression (first principal component of the gene expression matrix) is plotted across developmental stages for four species, with colors corresponding to module assignments. Core transcription factors were identified based on co-expression dynamics and intersection with a curated human transcription factor list. **(B)** Genomic span and transcript length distributions for HLD, background, and conserved genes, stratified into 10 expression-level bins (independent of cell type). Gene length distributions are visualized, with p-values calculated using a corrected Kolmogorov–Smirnov test. **(C)** Human Accelerated Region (HAR) interactions, derived from published Hi-C data (see Methods), and epigenetic modifications from human and mouse ENCODE projects. Heatmaps and density plots depict regions 3 kb upstream and downstream of the gene body, as well as the gene body itself. Color scales are normalized to the global maximum and minimum values across all heatmaps.

Among the HLD genes, we observed a strong bias toward upregulation in the human lineage, suggesting a directional asymmetry in gene expression evolution. This trend points to a widespread enhancement of transcriptional output for specific genes during human neocortical development. To uncover the genomic features associated with this upregulation, we examined the sequence characteristics of human-upregulated genes and identified two prominent trends: these genes tend to have lower GC content and longer gene length compared to others in the human genome (Fig. 3B and Supplementary Fig. 14). Notably, a clear negative correlation between genomic GC content and expression level was observed only in humans, with no such relationship detected in macaque, mouse, or ferret. Furthermore, in humans, gene expression level showed a strong positive correlation with gene length—a pattern absent in the other species. To assess whether these relationships are unique to the human-upregulated gene set, we analyzed a control group of “conserved” genes, defined as those exhibiting minimal interspecies differences across all three axes. These conserved genes generally displayed moderately high expression levels but showed no significant correlation with gene length. Importantly, the length-dependent expression trend in humans was consistently observed across all cortical cell types (Supplementary Fig. 15), suggesting that this is a general feature of the developing neocortex rather than a cell-type-specific phenomenon. To further investigate whether gene length is a shared property among all categories of human-distinctive genes, we compared the gene lengths of HLD, HTD, and HDD classes (Supplementary Fig. 16). Strikingly, only HLD genes were enriched for longer length, whereas HTD and HDD genes did not show this trend. These findings indicate that the association between gene length and elevated expression is specific to the human-upregulated HLD genes and may reflect an evolutionary remodeling of transcriptional regulation unique to the human lineage.

To determine whether gene elongation might have occurred in the human evolutionary lineage, we assessed gene length across species (Fig.3B). These genes were already significantly longer than average in macaque, mouse, and ferret. Thus, gene length was not a derived feature of the human lineage, but rather a pre-existing attribute that was later coupled with transcriptional upregulation in humans. This implies that a subset of already long genes underwent increased expression in the course of human brain evolution. To further probe this relationship, we analyzed the length of spliced, mature transcripts and found no correlation with expression level (Fig.3C). This indicates that the relationship between gene length and expression is primarily driven by intronic, rather than exonic, sequences. Taken together, our findings suggest that genes with unusually long introns are preferentially upregulated in the human neocortex, pointing to a potential regulatory role for these non-coding regions.

We next asked whether these long genes exhibit any other distinctive features beyond their genomic length. Given the extensive size of their introns, we hypothesized that regulatory elements, such as enhancers, might be embedded within these non-coding sequences. To test this, we analyzed epigenomic profiles of human neural stem cells, neurons, and astrocytes from the ENCODE project, but found no consistent enrichment of classical histone modification marks that would clearly distinguish these genes (Fig.3D). However, because we sought to understand mechanisms underlying human-specific transcriptional upregulation, we turned to human accelerated regions (HARs), most of which have been reported to harbor enhancer activity to regulate expression of HLD genes^41,49,50^. Strikingly, we found that genomic regions exhibiting physical interactions with HARs were significantly enriched within or near the gene bodies of these genes (Fig.3E). Among the 267 HLD genes identified as significantly upregulated or downregulated in humans, 117 genes showed evidence of interaction with HAR-associated elements. This enrichment was specific to HLD genes; neither HTD genes (29 / 187) nor HDD genes (24 / 222) showed similar association with HARs (background as 1324 / 10684 genes). This suggests that HARs may have contributed specifically to the transcriptional upregulation of certain long, intron-rich genes during human evolution.

In summary, the vast majority of HLD genes exhibit elevated expression in human cortical cells compared to those of other species. This gene set is notably enriched for genes involved in synaptic function and intercellular communication, despite the fact that the analyzed cells are largely derived from developmental stages preceding the onset of synaptogenesis. These genes tend to be disproportionately long, particularly with respect to intronic regions, and show a high frequency of physical interactions with HARs. Together, these findings support a model in which evolutionary changes in HAR-associated regulatory elements have driven increased transcription of selected long genes in the developing human neocortex, potentially contributing to the emergence of human-specific features in cortical architecture and function.

### Human distinctive genes with altered temporal expression trend

To investigate the evolution of gene expression temporality during neocortical development, we compared the temporal progression of transcriptomic profiles from early to late developmental stages across species. Specifically, we focused on a set of 206 HTD genes that showed consistent human-specific divergence in temporal expression trends, i.e., genes whose expression dynamics over developmental time differed markedly in humans compared to macaque, mouse, and ferret. In all three pairwise comparisons, these 206 genes displayed a striking and consistent pattern: whereas their expression levels increased over time in non-human species, they peaked early and declined thereafter in humans (Fig.4A-C and Supplementary Fig.17). This inversion of temporal dynamics appeared unique to the human lineage, as comparisons among non-human species revealed no such divergence in the expression trajectories of the same orthologs (Fig.4D-F). Inspection of the gene list revealed an overrepresentation of functionally coherent gene categories (Fig.4G-J and Supplementary Fig.13). Gene ontology analysis showed significant enrichment for genes involved in oxidative phosphorylation (e.g., NDUFS5, NDUFA6, NDUFA2, NDUFB4, NDUFB2, NDUFA13) and components of the cytosolic large ribosomal subunit (e.g., RPL36AL, RPL21, RPL17, RPL7, RPL32, RPL29). The temporal reprogramming of oxidative phosphorylation–related genes was particularly notable: while their expression levels increased progressively from early to late stages in macaque, mouse, and ferret, they were highest at the earliest stage in humans and declined steadily over time—a clear inversion of the typical developmental trend (Fig.4G and Supplementary Fig.18). To validate these observations, we examined independently generated bulk RNA-seq data from human and chimpanzee cortical organoids (Fig.4H). Consistent with our *in vivo* findings, human organoids exhibited high early expression and subsequent downregulation of these genes. In contrast, chimpanzee organoids showed a flatter expression profile, lacking the pronounced early peak seen in humans. These results support the notion that this temporal shift is a derived, human-specific feature. A similar but subtler pattern was observed for ribosomal genes, which also peaked early in humans but remained relatively flat in other species (Fig. 4I-J and Supplementary Fig.18). Interestingly, this inverted temporal pattern was broadly consistent across cell types and differentiation lineages, suggesting that it reflects a global regulatory program rather than cell type–specific modulation (Fig.4K).

**Figure 4.**
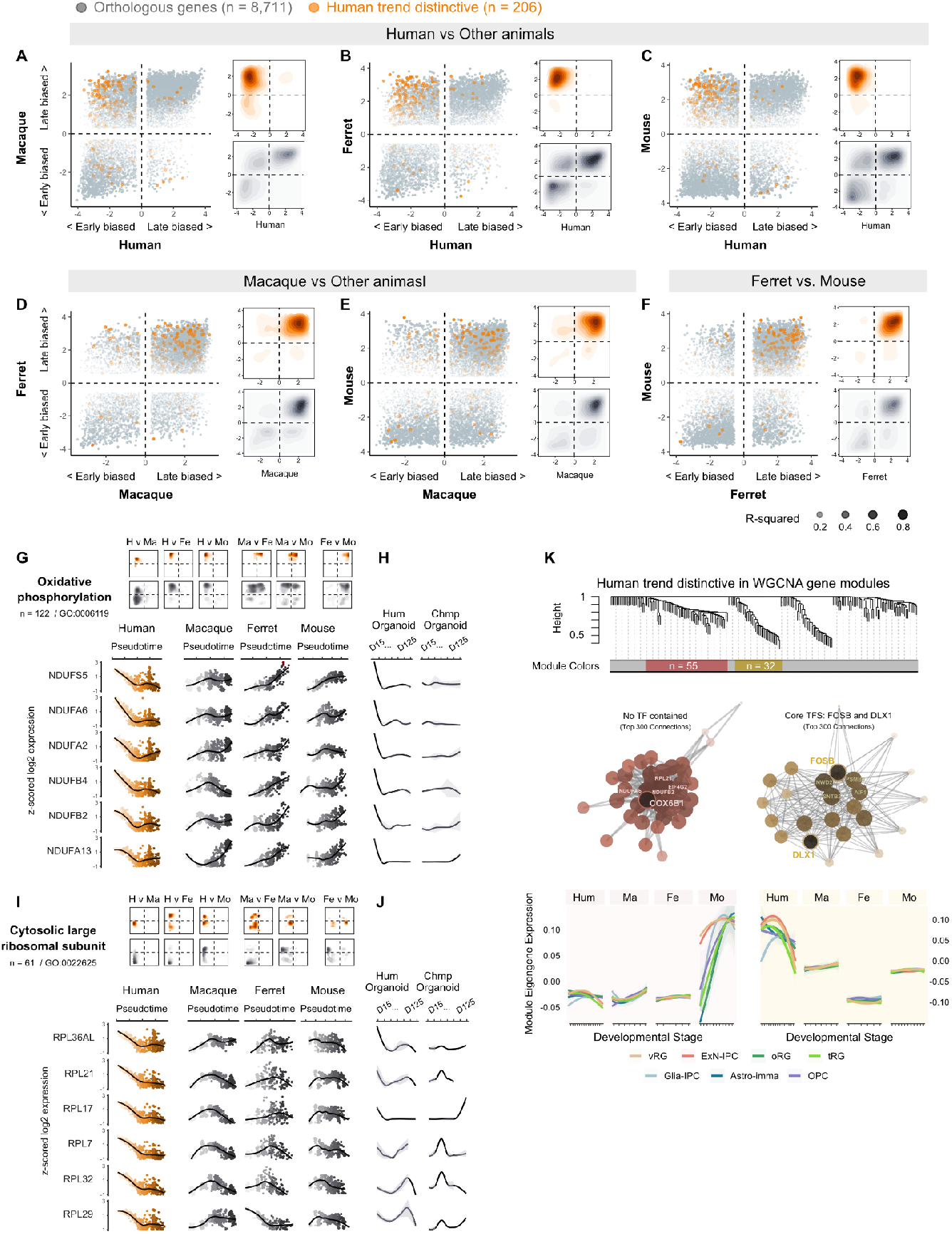
Human Trend Distinctive (HTD) Genes Exhibit Opposite Expression Trends. **(A–C)** Comparison of gene expression trends in human versus macaque, ferret, and mouse across developmental stages. Linear regression coefficients for each species are calculated, and gene covariation is plotted in a 2D space. Kernel-estimated density distributions are shown on the right. **(D–E)** Comparison of gene expression trends in macaque versus ferret and mouse, following the methodology described in (A–C). **(F)** Comparison of gene expression trends in ferret versus mouse. Notably, human HTD genes exhibit a unique early-high, late-low expression trend, which is opposite to trends observed in macaque, ferret, and mouse. **(G–H)** Analysis of genes associated with Oxidative Phosphorylation GO term (n=112). Density distributions of these genes in species comparisons are shown, alongside their expression dynamics across pseudotime in each species. Color gradient (light to dark) indicates actual sampling stages. Expression dynamics in human and chimpanzee cerebral organoids are presented on the right. Expression values are z-scored and centered. **(I–J)** Analysis of genes involved in the Cytosolic Large Ribosomal Subunit (n=61), following the same methodology as in (G–H). **(K)** Weighted Gene Co-expression Network Analysis (WGCNA) of HTD genes identifies two modules. Co-expression networks and module eigengene expression across developmental stages are shown, with corresponding module-specific trends highlighted.

To further dissect the regulatory basis of this phenomenon, we performed WGCNA on the 206 HTD genes (Fig.4K). This yielded two major co-expression modules. One “red” module consisted primarily of genes related to oxidative phosphorylation and ribosomal proteins (e.g., COX6B1, NDUFA6, RPL21), but did not contain identifiable transcription factors. The second “yellow” module included transcriptional regulators such as FOSB and DLX1, which occupied central positions in the network, suggesting a potential upstream role in orchestrating the coordinated temporal downregulation observed in humans.

In summary, we identified a cohort of genes whose expression dynamics were uniquely reprogrammed in the human neocortex, characterized by early high expression followed by a gradual decline—opposite to the patterns observed in non-human mammals. These genes are enriched for roles in aerobic metabolism and protein synthesis, and their coordinated temporal shift may be regulated by transcription factors such as FOSB and DLX1. This human-specific alteration in gene expression timing could reflect an evolutionary adaptation in the metabolic and translational landscape of early neurodevelopment.

### Human-distinctive genes with altered differentiation lineage specificity

To explore human-specific alterations in the cellular trajectories of gene expression during neocortical development, we analyzed 227 HDD genes that exhibited changes in lineage specificity unique to humans. The predicted cell type-to-cell type lineage trajectories demonstrate the expected curves of marker gene expressions (Fig.5A). Gene ontology (GO) analysis revealed a striking enrichment of genes associated with cilia and axonemal microtubule function, including CFAP46, DRC1, and DAW1 (Supplementary Fig.13). The cilia-related genes found in HDD show notable human distinctive expression dynamics in the trajectories containing oRG progenitor (Fig.5B).

In the developing neocortex, primary cilia are most prominently localized at the apical ventricular surface, where neural progenitors such as vRGs and tRGs reside (Fig. 5C). Consistent with this localization, cilia-related genes in macaque, mouse, and ferret showed their highest expression levels along the differentiation trajectory from vRG to tRG (Fig. 5B, D). In humans, however, this spatial expression pattern was markedly different. Cilia-related genes exhibited elevated expression not in the vRG-to-tRG lineage, but instead along trajectories involving oRGs (Fig.5B). Specifically, expression levels peaked during the late stages of the vRG-to-oRG transition and the early stages of oRG-to-neuron or oRG-to-glia differentiation. These findings suggest a human-specific redistribution of ciliary gene expression from apical vRG progenitors to basally located oRGs. In other words, genes that are typically confined to ventricular zone progenitors in other species have become preferentially expressed in oRGs in humans, indicating a spatial reallocation of ciliary gene function during cortical development.

**Figure 5.**
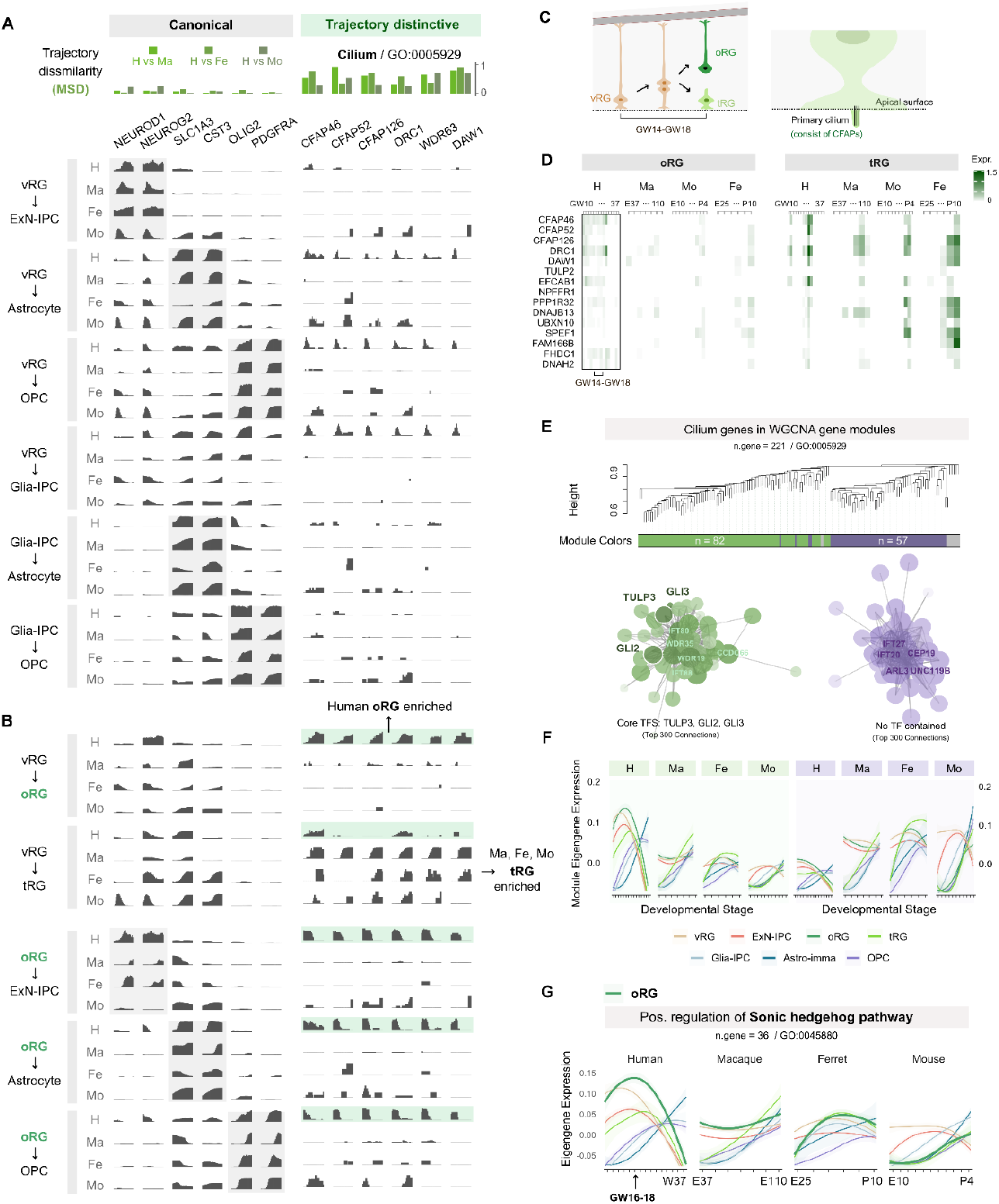
Distinctive Trajectories Bias of Cilium-Associated Genes in Human oRG trajectory. **(A–B)** Gene expression along inferred differentiation trajectories. Six canonical fate-driving or marker genes are listed on the left, and six representative cilium-associated genes from the Human Differentiation Trajectory (HDD) gene set are listed on the right. The x-axis represents trajectory progression, and the y-axis denotes expression levels. Notably, cilium-associated genes in humans are preferentially expressed in the outer radial glia (oRG) lineage, unlike in other species where expression is predominant in truncated radial glia (tRG). **(C)** Schematic illustrating the divergence of ventricular radial glia (vRG) into oRG (with basal fibers) and tRG (with apical fibers, retained at the apical surface) during mid-neurogenesis. **(D)** Heatmap depicting cilium-associated gene expression in oRG and tRG across developmental stages in four mammalian species. **(E–F)** Weighted Gene Co-expression Network Analysis (WGCNA) identifies two modules within cilium-associated genes. The green module exhibits a pattern similar to the HDD cilium genes, with the most prominent expression in human oRG compared to tRG. **(G)** Expression dynamics of genes involved in the positive regulation of the Sonic Hedgehog (SHH) signaling pathway. In humans, the expression peak coincides with oRG emergence and the peak of cilium-associated gene expression, the latter being an upstream regulator of the SHH signaling pathway.

Given that primary cilia act as signaling hubs by concentrating key receptors, we next examined the functional implications of this shift. Many of the human-distinctive ciliary genes encode components of the intraflagellar transport machinery, such as microtubule-binding proteins and motor proteins required for trafficking signaling molecules within the cilium. Proper trafficking is essential for the reception and transduction of extracellular cues, particularly Sonic Hedgehog (SHH) signaling, which has been shown to promote oRG proliferation and enhance neuronal output during cortical development^51,52^. GO analysis further revealed enrichment of genes involved in “Signaling receptor binding,” including BMP4, ITGB1BP2, and IGFBP6 (Supplementary Fig.13). To explore the broader transcriptional impact, we analyzed the expression profiles of all “cilium” (GO:0005929) genes across the four species during cortical development. WGCNA identified two major modules: one containing SHH signaling effectors *GLI2* and *GLI3* (green module), which showed marked upregulation in human oRGs; and another module whose genes were expressed at similar or lower levels in human oRGs compared to other species (Fig.5EF). Additionally, comparative analysis of genes assigned to the “SHH signaling pathway” (GO:0045880) revealed consistently higher expression in human oRGs than in the corresponding cell type of macaque, mouse, or ferret (Fig.5G).

Taken together, these findings suggest that the spatial reorganization of cilia-related gene expression to the oRG lineage represents a human-specific adaptation in progenitor signaling architecture. This reprogramming may underlie enhanced oRG activity and expanded neurogenic potential in the developing human neocortex.

### Disease relevance of human-specific gene expression changes

Through comparative analysis of gene expression dynamics across species, we identified three distinct groups of genes exhibiting human-specific features. Given that these gene sets were enriched in distinct functional categories, it is likely that each contributes to different biological consequences in human brain development and evolution. To assess whether these human-specific expression changes might be linked to disease susceptibility, we systematically examined the relationship between each gene group and a wide range of brain-related disorders (Fig.6A). We compiled curated gene lists associated with neurodevelopmental disorders, brain size abnormalities, miscarriage and congenital malformations, as well as brain tumors^53–77^ (Supplementary Table 3). Then, we examined the positions of the genes associated with a given disease in “Gene Expression Dissimilarity Space” and calculated the centroid position of the gene set, which indicates the general trend of the evolutionary change in the expression of the genes associated with a certain disease (Fig.6A and Supplementary Fig.19). A deviation vector was computed from the centroid of all homologs ( n = 10,683) to that of each disease gene set, highlighting expression shifts specifically attributable to the disease, beyond inherent species differences. This vector was further decomposed into its x, y, and z components to provide a detailed characterization of the evolutionary alterations in gene expression. To address the variability in gene set sizes across different diseases, centroids were employed, and permutation tests using random gene sets of equivalent size were conducted to ensure a fair and unbiased evaluation (Fig.6A).

Among the three classes of gene expression evolution, those with elevated expression in humans showed the strongest enrichment for disease-associated genes (Fig6B). Notably, genes implicated in developmental delay (DD) (Fig.6C), autism spectrum disorder (ASD) (Fig.6D), megalencephaly (MEG) (Fig.6E), and Structural abnormality (Stur) (Fig.6F) were significantly overrepresented in this group. Glioblastoma multiforme (GBM), a malignant brain tumor, also showed marked enrichment for genes with elevated expression, consistent with prior reports highlighting shared genetic risk between neurodevelopmental disorders and GBM (Fig.6G). In contrast, genes with reduced expression in humans were enriched for Alzheimer’s disease (AD)–associated genes, indicating a possible inverse pattern of evolutionary expression shift (Fig.6H). Furthermore, neuroblastoma and medulloblastoma were more frequently associated with genes that showed minimal changes in temporal trends or lineage specificity, suggesting these diseases may involve evolutionarily conserved pathways.

When examining individual genes associated with ASD, DD, MEG, and AD, the trends became even more pronounced (Fig.7A). Among the three classes of human distinctive genes, only HLD genes were significantly enriched in disease-associated gene sets, suggesting that these genes exhibit cell type–independent upregulation during human cortical development. Comparative analysis using cortical organoids confirmed that these genes are expressed at significantly higher levels in humans than in chimpanzees, further supporting the notion that this upregulation is a feature specific to the human evolutionary lineage. Genomic analysis further revealed a significant enrichment of HAR-associated elements within the loci of these genes, suggesting potential regulatory contributions. Notably, several HLD genes, including AUTS2 and NLGN1, which are well-established regulators of synaptic development and are implicated in various neurodevelopmental disorders—showed substantially elevated expression in humans (Fig. 7B). This species-specific expression pattern was corroborated by immunohistochemical analysis, which confirmed high levels of protein expression in human cortical organoids, but not in embryonic mouse cortex (Fig. 7C). More specifically, cell type specific expression patterns of AUTS2 and NLGN1 are confirmed by immunohistochemistry on human cortical organoids; AUTS2 is most prominently detected in ExN-IPC, which is the cells immediately above the ventricular zone-like region, and NLGN1 is broadly detected in progenitor subtypes, in particular vRG in the early stages, which is also confirmed experimentally.

**Figure 6.**
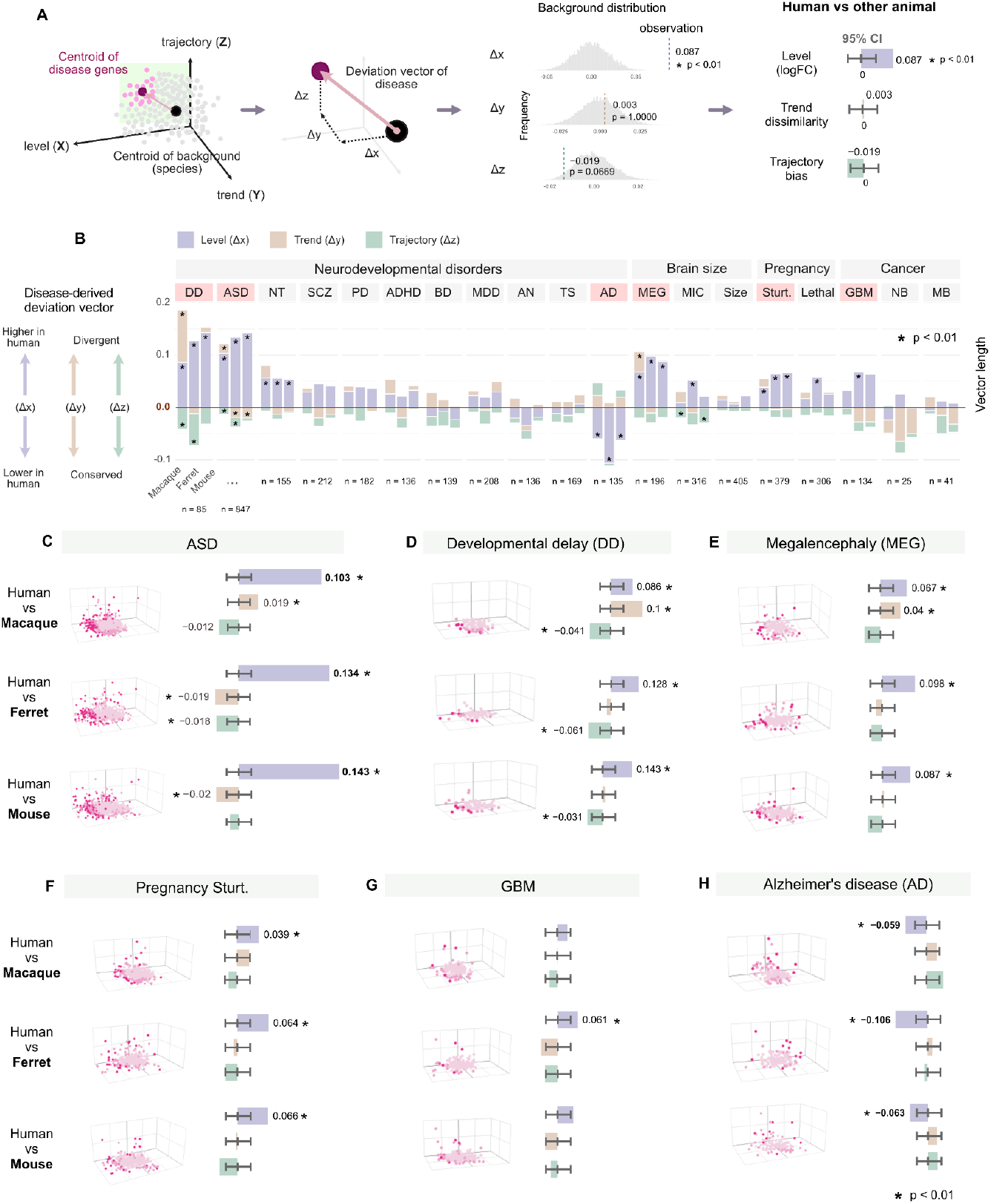
Disease-Associated Gene Expression Deviation Vectors in Cross-Species Divergence Space. **(A)** Schematic illustrating the identification of disease-derived deviation vectors. A vector is defined from the centroid of the background gene set (all genes) to the centroid of disease-associated gene sets, decomposed into three components: Δx (level), Δy (trend), and Δz (trajectory), representing the magnitude, trend, and trajectory deviations, respectively. Statistical significance is assessed by comparing the observed deviation vector to 10,000 randomly sampled vectors of equal gene count, calculating the proportion of random vectors with greater deviation. The 95% confidence interval (CI) of random vector deviations is computed and marked with ticks at corresponding positions. **(B)** Summary of deviation vectors in divergence space for 19 diseases, with disease-associated gene sets sourced from public GWAS, MAGMA, and curated databases (see references). Diseases include Alzheimer’s disease (AD), anorexia nervosa (AN), autism spectrum disorder (ASD), bipolar disorder (BD), major depressive disorder (MDD), Parkinson’s disease (PD), schizophrenia (SCZ), structural abnormality (Sturt.), glioblastoma (GBM), neuroblastoma (NB), medulloblastoma (MB), megalencephaly (MEG), and microcephaly (MIG). **(C–H)** Representative disease deviation vectors from the summary. Notably, ASD exhibits the most pronounced Δx (elevated in human), while AD-associated genes show a unique negative Δx deviation (reduced in human).

**Figure 7.**
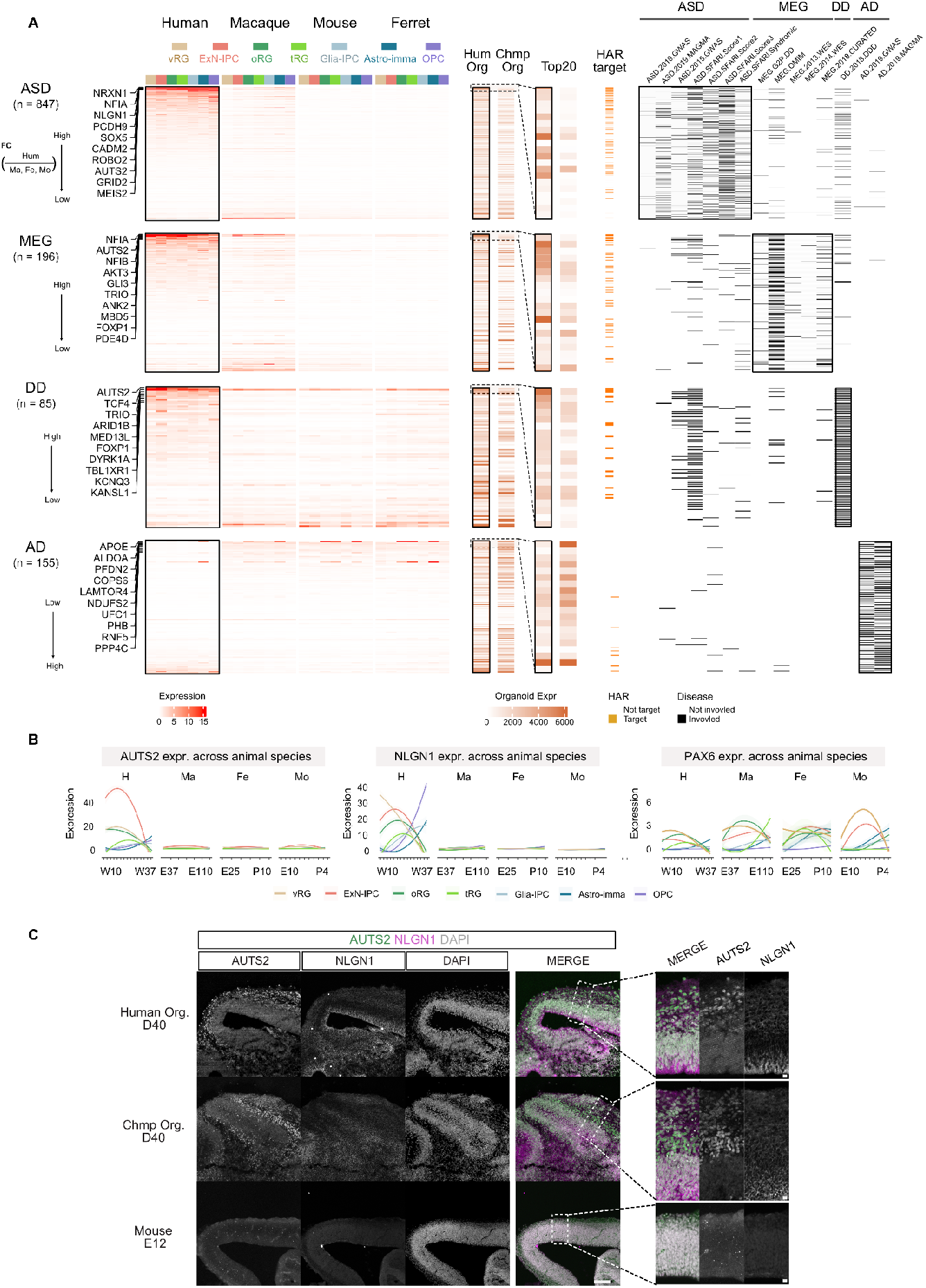
ASD-, DD-, MEG-, and AD-Associated Genes Expression Across Cell Types and Mammalian Species. **(A)** Heatmap depicting the expression of genes associated with autism spectrum disorder (ASD), developmental delay (DD), megalencephaly (MEG), and Alzheimer’s disease (AD) across seven cell types in four mammalian species (human, macaque, mouse, ferret) using single-cell RNA-seq data, validated with human and chimpanzee cortical organoids. Gene expression is modulated by interactions with Human Accelerated Regions (HARs). Rows are sorted by the log fold change of human expression relative to the average expression of the other three species. **(B)** Expression profiles of AUTS2 and NLGN1, selected from the gene set, plotted across developmental stages in the four species. PAX6, a canonical marker gene, is included as a negative control. **(C)** Immunostaining of AUTS2 and NLGN1 in day 40 (D40) human and chimpanzee cortical organoids and embryonic day 12 (E12) mouse neocortex. Scale bar: 100 μm (left panel); 10 μm (enlarged panel).

Collectively, these findings indicate that human-specific changes in gene expression, especially those that increased transcriptional output, may underlie both the evolutionary expansion of the human cortex and its vulnerability to specific developmental disorders, including the brain size abnormality, and brain tumors. HAR-associated regulation emerges as a plausible mechanism driving these dual outcomes.

## Discussion

To identify genes involved in the evolutionary expansion and complexification of the human neocortex, we conducted a comparative transcriptomic analysis across humans and three other mammalian species. Focusing specifically on the dynamic progression of neural progenitors into differentiated cortical cell types during development, we extracted only proliferative cell populations to enable detailed comparison of gene expression patterns across species. Throughout mammalian evolution leading to humans, we observed a variety of gene expression changes that could be broadly categorized into three distinct groups. The first group comprises genes that exhibit changes in overall expression levels (“heterometric” changes). The second includes genes showing altered temporal expression trend, i.e., “heterochronic” changes. The third group consists of genes that underwent shifts in lineage specificity, expressing in different developmental trajectories (“heterotopic” changes). These categories are not mutually exclusive, and individual genes may display multiple types of expression divergence. We defined human-distinctive genes as those ranking in the top 5% in at least one of the three axes of expression divergence. Each group of human distinctive genes is enriched for distinct functional categories and disease relevance, suggesting that they may have been shaped by different evolutionary constraints and selective pressures.

### Rationale for classifying gene expression changes along three axes

Gene expression can evolve in various ways throughout the course of biological evolution. In this study, we employed a unified framework to visualize the diversity of gene expression changes by quantifying them along three well-established axes: expression magnitude (“heterometric”), temporal dynamics (“heterochronic”), and lineage specificity (“heterotopic”). These axes capture distinct aspects of transcriptional regulation and have been widely used in developmental and evolutionary biology^43^. Our analysis revealed that these three axes are largely independent (Fig.2). Among the genes exhibiting the strongest divergence in the human lineage, the majority showed pronounced changes along only one of the three axes. This finding suggests that evolutionary changes in gene expression tend to proceed along specific, non-overlapping trajectories—an observation consistent with previous studies.

By focusing on these three dimensions of expression change and extracting genes that exhibited the most pronounced divergence across pairwise comparisons between humans and three other species, we were able to systematically isolate genes that are evolutionarily conserved among non-human species but have undergone human-specific expression changes. This approach effectively narrows down candidate genes that may have contributed to uniquely human features.

Taken together, our framework for cross-species transcriptomic comparison based on these three axes provides a powerful strategy for identifying evolutionary shifts in gene regulation. While this study focused on cortical development, the same methodology could be broadly applicable to other biological contexts with temporal progression, including organogenesis and aging. Given single-cell transcriptomic datasets sampled at multiple developmental time points across multiple species, this approach can yield high-confidence candidates for genes driving lineage-specific phenotypic innovations.

### HLD: human level distinctive genes showing significant changes in expression magnitudes

Among the three dimensions analyzed, genes exhibiting quantitative changes in expression levels (HLD genes) displayed striking consistency across the three species comparisons (human vs. macaque, mouse, and ferret) (Fig.3). This pattern suggests that while the expression levels of these genes have been relatively conserved throughout mammalian diversification, they underwent significant transcriptional shifts specifically along the human lineage. Notably, the vast majority of these genes showed increased expression in humans, whereas only a small subset exhibited decreased expression. Although a small subset of downregulated HLD genes have a co-expression network structure with ZBTB12 as a potential core transcriptional regulator, the underlying mechanisms driving their reduced expression remain unclear. In contrast, the upregulated genes tend to have significantly longer gene bodies than the genomic average. This length bias is observed even in the non-human species, indicating that gene length itself is not the consequence of expression upregulation but rather a precondition enabling such upregulation in the human lineage. Interestingly, this trend does not apply to the length of mature, spliced transcripts, implying that intron length—rather than exon content—may play a role in modulating transcriptional output. Supporting this hypothesis, the intronic regions of these long genes are highly enriched for sequences that interact with HARs. These observations suggest a model in which evolutionary changes in HARs, possibly through chromatin-mediated mechanisms, enhanced transcriptional activity of their associated genes. This scenario parallels the previously described case of *Fzd8*, whose expression is elevated by the enhancer HARE5^39,40^. Several other genes previously linked to HAR-mediated regulatory evolution during cortical development, such as AUTS2 and NPAS^37,38,41,42,50,78,79^, were also identified in our analysis, underscoring the consistency of our findings with earlier reports.

Intriguingly, many of the HLD genes that exhibit upregulated expression in humans are typically associated with postnatal neurodevelopmental processes—such as glutamatergic signaling and synapse formation—in non-human species. However, these genes are already expressed at high levels during the earlier neurogenic phase of cortical development in humans. Immunohistochemical analysis of human cortical organoids confirmed that key genes such as AUTS2 and NLGN1 are expressed across a broad range of progenitor and differentiated cell types, including vRGs. Given that many of these genes are strongly implicated in neurodevelopmental disorders such as ASD and DD, their precocious and elevated expression during early corticogenesis may play a previously unrecognized role in regulating neural progenitor behavior and neuronal output. For instance, neural progenitor proliferation and differentiation are known to be influenced by intrinsic electrophysiological properties, such as membrane potential changes and calcium transients^80–88^, processes that may be modulated by the early expression of genes involved in intercellular signaling pathways like glutamatergic transmission. These observations highlight the need for future functional studies using human brain organoids and complementary models to elucidate the mechanistic significance of this human-specific regulatory shift.

### HTD: human trend distinctive genes showing significant changes in temporal dynamics of expression

Heterochronic changes in gene expression have long been implicated in the evolutionary expansion of the human brain. In this study, we focused not on absolute or physical developmental time, but on *relative* timing within a shared developmental window spanning from the onset of neurogenesis to the beginning of gliogenesis. To this end, we applied pseudotime analysis to identify genes that exhibit distinct temporal expression profiles in humans^89^. Importantly, this approach does not address heterochronicity in the classical sense of delayed neurodevelopment (i.e., neoteny), in which the onset of developmental events is globally postponed in humans^1,2,4,8,17^. Rather, our analysis aims to uncover genes that, despite being involved in the same overall sequence of developmental transitions across species, are expressed at divergent timepoints *within* that sequence in humans alone.

This analysis revealed a consistent trend: genes expressed during the late stages of cortical development in non-human species tend to be upregulated earlier in humans. This pattern was not observed in comparisons between non-human species (e.g., macaque vs. ferret or ferret vs. mouse), but emerged consistently across all human vs non-human comparisons. These findings indicate that a shift in the timing of gene expression, in which specific transcripts are precociously activated in the human lineage, represents a human-specific evolutionary feature.

Strikingly, the genes exhibiting human-specific shifts in temporal expression dynamics were enriched for components involved in mitochondrial oxidative phosphorylation and the translational machinery, including ribosomal proteins. These two fundamental cellular processes—mitochondrial metabolism and protein synthesis—have repeatedly been implicated in regulating developmental timing and explaining interspecies differences in developmental speed ^90–93^. It is well established that slower-developing species, typically humans, generally exhibit globally reduced mitochondrial and translational activity compared to faster-developing species like mice. This phenomenon has been documented in various developmental contexts, including somite formation^94–96^, spinal neuron specification^97^, and postmitotic neuronal maturation^98^. However, our data provide a more nuanced view. While overall mitochondrial and translational activity remain low in humans, we observed a transient upregulation of mitochondrial and ribosomal genes during the earliest phase of cortical neurogenesis. This pattern suggests that early neural progenitors may undergo a short period of metabolic acceleration, which is later attenuated as development progresses. This observation is consistent with previous findings that mitochondrial fusion—known to promote oxidative phosphorylation—is prominently observed in undifferentiated neural progenitors and contributes to the maintenance and proliferative capacity of these cells^90,99–102^. Such a temporally restricted enhancement of metabolic activity may act as a regulatory mechanism to promote early amplification of progenitor pools specifically in human corticogenesis. These findings raise the hypothesis that precise modulation of mitochondrial and translational functions at early developmental stages could be a key determinant of species-specific neurogenic trajectories. This model could be experimentally tested by manipulating these pathways in human and non-human primate cortical organoids during the initial phase of cortical development.

Co-expression network analysis further supports the idea of transcriptional regulation underlying these shifts. A subset of the early-expressed genes shared dynamic expression patterns with transcription factors such as FOSB and DLX1, which occupy central nodes in the regulatory network structure. These findings suggest that such transcription factors may play pivotal roles in orchestrating early developmental gene expression programs that are uniquely regulated in the human lineage. It is worth testing the importance of these transcription factors in the human-specific temporal dynamics of mitochondrial and ribosomal genes by knocking them out in human cortical organoids.

### HDD: human differentiation trajectory distinctive genes showing altered expression preference on cell differentiation lineages

To investigate heterotopic changes in gene expression during human neocortical development, we systematically identified genes that exhibited species-specific alterations in lineage trajectories. Notably, a large subset of these genes was functionally linked to primary cilia. This shift in expression may reflect either an evolutionary enhancement of ciliary structures in human oRGs or a repurposing of ciliary components for novel functions within these progenitors. In non-human species, cilia-related genes were predominantly expressed in vRGs and tRGs, both of which are anchored at the apical ventricular surface—a region classically associated with primary cilia^103^. In contrast, human cortical tissue exhibited robust expression of these same genes not only in apical progenitors but also in oRGs, which are located away from the ventricular surface. This indicates a human-specific spatial shift in the deployment of ciliary gene expression. Importantly, this shift in expression trajectory may act synergistically with other human-specific ciliary adaptations. We previously identified CROCCP2, a human-specific gene that regulates cilia length through interactions with IFT20 and mTOR signaling, ultimately promoting basal neurogenesis^26^. Together, these findings point to the central role of primary cilia in human-specific aspects of cortical development and underscore the importance of further mechanistic investigation into cilia-dependent processes in oRGs.

The precise status and function of primary cilia in basal progenitors detached from ventricular surface remain incompletely understood. Prior studies in mouse and chick spinal cord have reported a phenomenon known as apical abscission, wherein differentiating neurons sever their apical membrane along with the primary cilium from the apical surface, retaining only the basolateral membrane as they migrate^104^. These neurons then reassemble a new primary cilium during migration^105^. Although apical abscission has not yet been directly demonstrated in the delaminating cells in the developing cerebral cortex, it is known that cilia in vRGs retract and shorten significantly during mitosis^106^. One of the daughter cells that inherit the retracted cilium tends to remain as vRGs within the ventricular zone, while its sister cell is more likely to differentiate into neurons or ExN-IPCs and migrate basally. Additional studies in mouse have shown that delaminating Tbr2-GFP+ cells can re-establish cilia on their basolateral surfaces^107^. Ciliogenesis in these cells begins around the time of delamination from the ventricular surface, even before full separation occurs. However, these observations were made mostly in mice that possess minimal numbers of oRGs, and the precise dynamics of cilia during the transitions of vRG-to-oRG and oRG-to-more differentiated cell types in humans remain unexplored.

Future experiments should take advantage of fluorescent cilia markers such as Arl13b-GFP, introduced into human pluripotent stem cells via genome editing. This would enable real-time tracking of ciliary dynamics during oRG generation in cortical organoids. If mature cilia are indeed a defining feature of human oRGs, it becomes crucial to investigate whether they serve as platforms for critical signaling pathways. Supporting this possibility, our data—as well as prior studies^51^—indicate that Sonic Hedgehog (SHH) signaling is more strongly activated in human oRGs than in their counterparts from other species. Functional experiments using human brain organoids, in which cilia are fluorescently visualized and SHH pathway components such as SMO and PTCH are tracked, will be essential to determine whether these molecular structures confer species-specific signaling competence during cortical development.

### Disease-relevance of human distinct genes

Genes that exhibit pronounced expression changes along the human lineage are likely to contribute to brain development in uniquely human ways and may underlie the emergence of advanced cognitive functions, many of which are altered in neurodevelopmental and neuropsychiatric disorders. To explore this potential link, we systematically assessed whether genes previously implicated in brain disorders also show human-specific expression changes, and if so, along which of the three regulatory dimensions those changes occur. Our analysis revealed that the directionality and nature of these changes varied depending on the disease category. Genes associated with DD, ASD, and MEG showed a strong bias toward elevated expression levels in humans relative to non-human species. Considering that a hallmark of human brain evolution is the dramatic expansion in brain size, the upregulation of many MEG-associated genes in the human lineage is particularly noteworthy. Likewise, the extended developmental timeline and neotenic features that distinguish humans from other primates may help explain the elevated expression of DD-related genes in the human fetal brain. In line with previous findings^13,108^, we also observed a substantial overlap between MEG-associated genes and those implicated in glioblastoma (GBM), suggesting that both conditions may involve disruptions in shared regulatory mechanisms that govern the number of neural cells. This convergence highlights a core set of genes, including GLI3, AKT3, AUTS2, NFIA, NRXN1, and NLGN1, that are likely to play central roles in both human brain evolution and disease susceptibility.

Most of these genes exhibit increased expression in humans across a broad range of cell types, spanning from neural progenitors to differentiated neurons, rather than being confined to a specific lineage. Notably, many of these genes harbor multiple regions of interaction with HARs, suggesting that their transcriptional regulation is likely influenced by HAR-associated enhancer activity. This interpretation aligns with previous findings showing that biallelic mutations within HARs are frequently observed in individuals with ASD^109^. Together, these observations support the hypothesis that evolutionary mutations giving rise to HARs contributed directly to the transcriptional upregulation of disease-associated genes during human brain evolution.

In contrast, genes linked to AD displayed a markedly different trend, showing reduced expression in the human fetal brain relative to other species. While the causal relationship between neurodegeneration and fetal developmental processes remains unclear, this finding highlights the need for comparative studies focusing on postmortem adult brains to investigate species-specific features of brain aging^110–113^. Unlike fetal samples, which are challenging to obtain from non-human primates, adult postmortem brain tissue is more readily accessible in both humans and non-human species.

Therefore, comparative transcriptomic analyses akin to the present study may be feasible in the context of aging and could provide critical insights into the evolutionary mechanisms underlying human susceptibility to neurodegenerative diseases.

## Summary

Our comprehensive multi-species analysis reveals that human neocortical evolution has been shaped by diverse gene expression changes, not just in magnitude, but critically, in their temporal dynamics and differentiation lineage specificity. These distinct evolutionary modes, mediated in part by genomic features such as HARs, highlight a complex interplay between conserved developmental programs and human-specific adaptations. The identified human-distinctive genes, particularly those exhibiting altered expression levels, are significantly enriched for neurodevelopmental and oncogenic disorders, suggesting that the very mechanisms driving human brain expansion and complexity may also confer heightened vulnerability to disease. Future research will focus on the functional validation of these candidate genes and their regulatory networks, leveraging advanced organoid models and gene-editing technologies to dissect the precise molecular mechanisms underlying human-specific cortical development and its associated pathologies.

## Supplementary Figures

**Figure S1.**
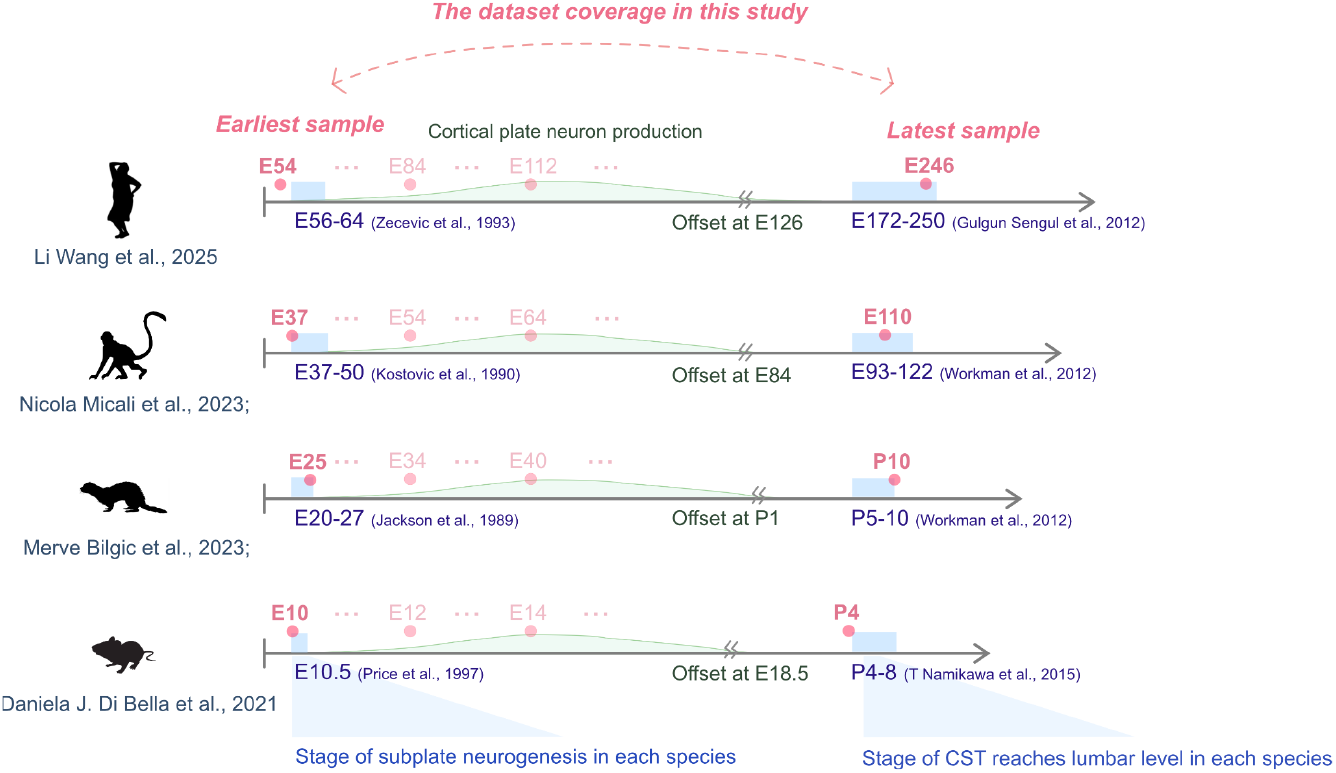
The dataset and developmental stage coverage in this study. We curated developmental stages from a public single-cell RNA sequencing database, using the emergence of subplate neurons as the starting point. The endpoint was defined by the arrival of the corticospinal tract at the lumbar level in each species, serving as the “outgroup” to align brain development timing across species. This timeframe encompasses the entire neurogenesis period and approximately the first half of gliogenesis, aligning with our aim to document and quantify neurodevelopment across multiple mammalian species for comparative analysis.

**Figure S2:**
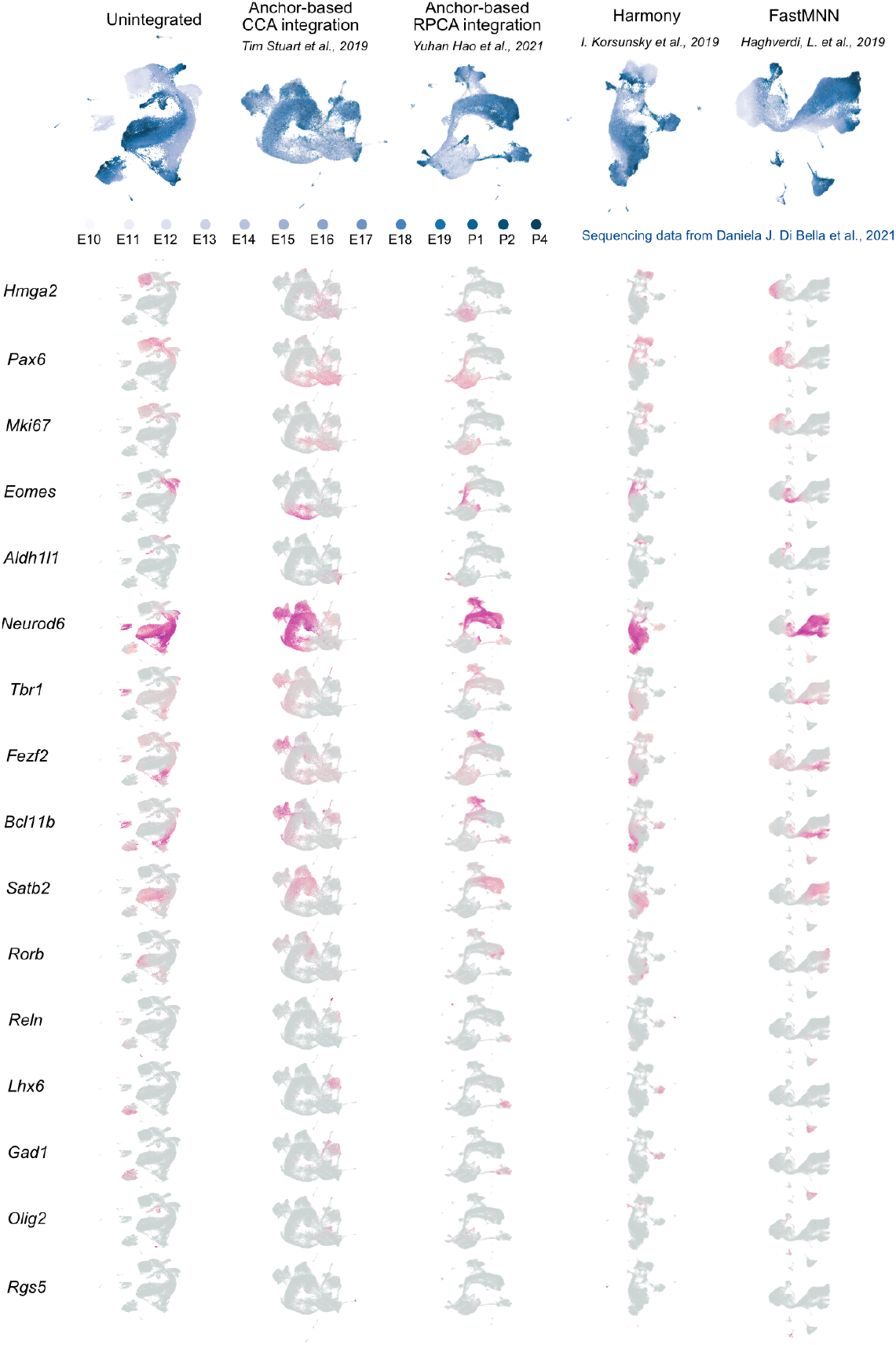
Categorization of Progenitor Cell Types in the Developing Mouse Cerebral Cortex. Five integration methods were applied to combine single-cell RNA-seq data from multiple developmental stages into a unified dataset, using canonical marker genes defined in this study. In the UMAP embedding space, cells are color-coded from light to dark blue to represent progression from early to late developmental stages. Expression patterns of canonical marker genes are visualized below the UMAP plots.

**Figure S3:**
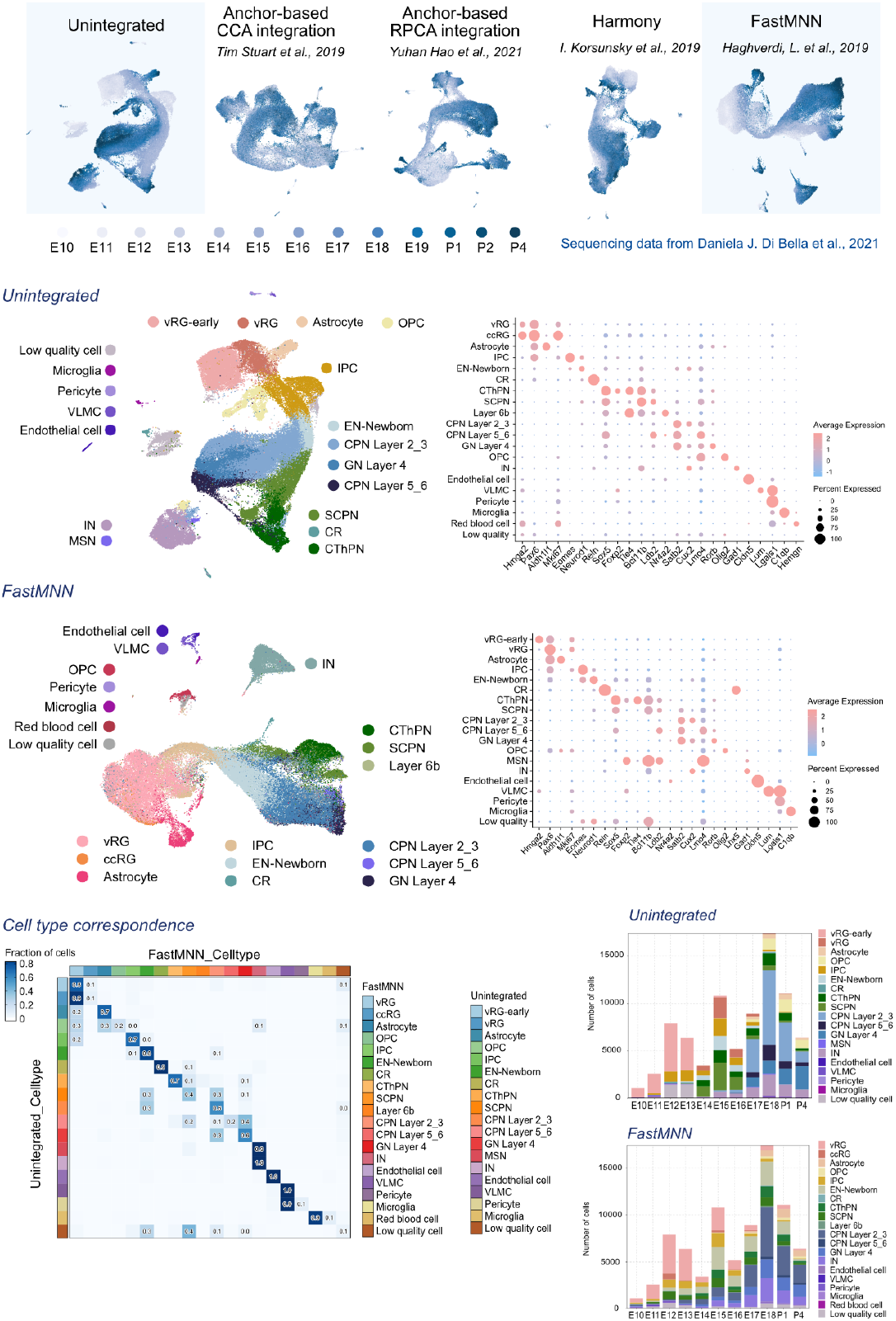
Cross-Validation of Cell Type Annotations in the Developing Mouse Cerebral Cortex. Unintegrated (SCT from Seurat) and FastMNN methods were selected and assessed for marker expression fidelity. Correspondence of cell type assignments across integration methods was calculated and visualized along developmental stages. Based on consistent marker expression, the unintegrated method (default regression provided by Seurat) was chosen for cell type annotation in the mouse dataset.

**Figure S4:**
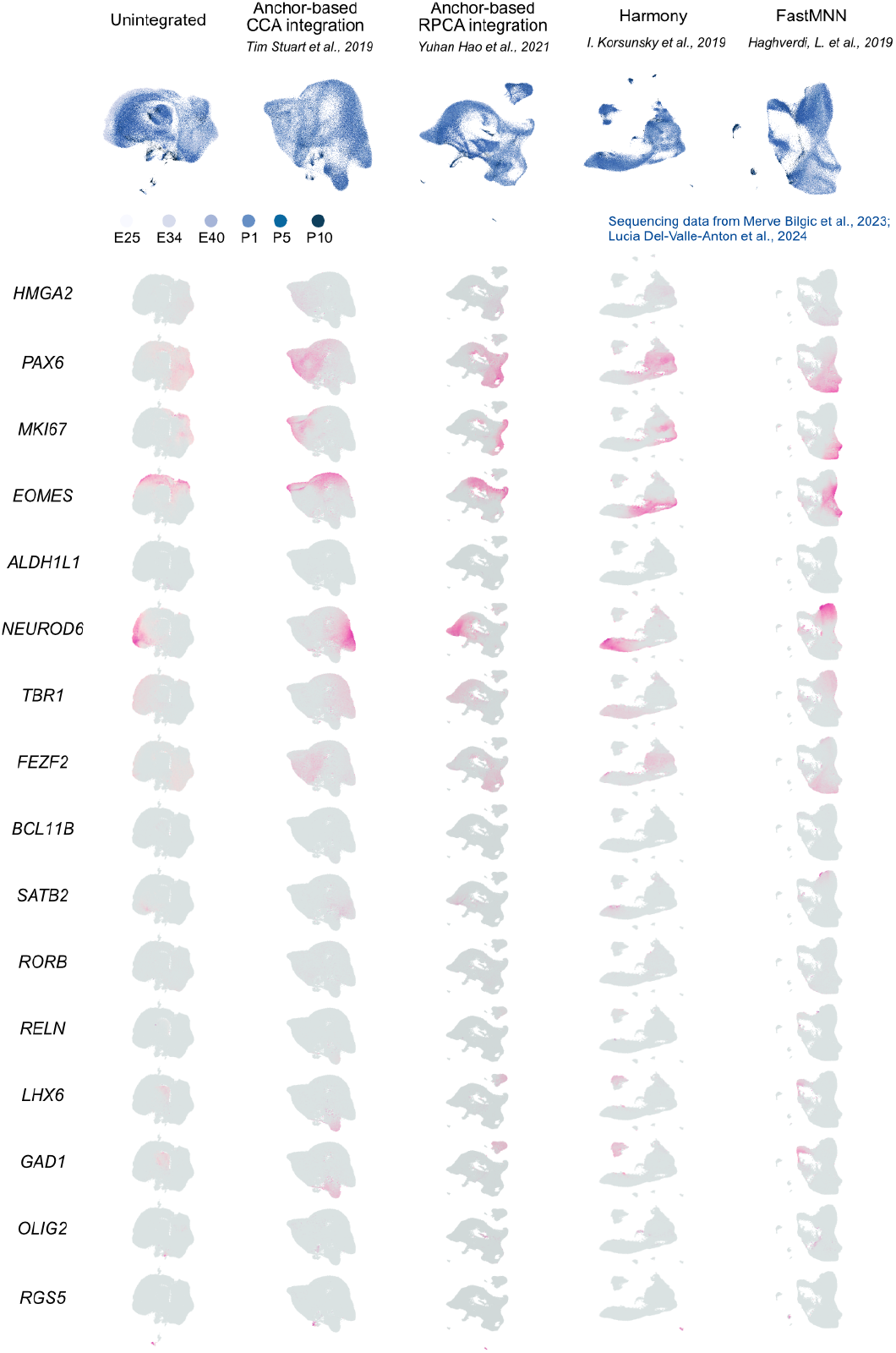
Categorization of Progenitor Cell Types in the Developing Ferret Cerebral Cortex. Five integration methods were applied to combine single-cell RNA-seq data from multiple developmental stages into a unified dataset, using canonical marker genes defined in this study. In the UMAP embedding space, cells are color-coded from light to dark blue to represent progression from early to late developmental stages. Expression patterns of canonical marker genes are visualized below the UMAP plots.

**Figure S5:**
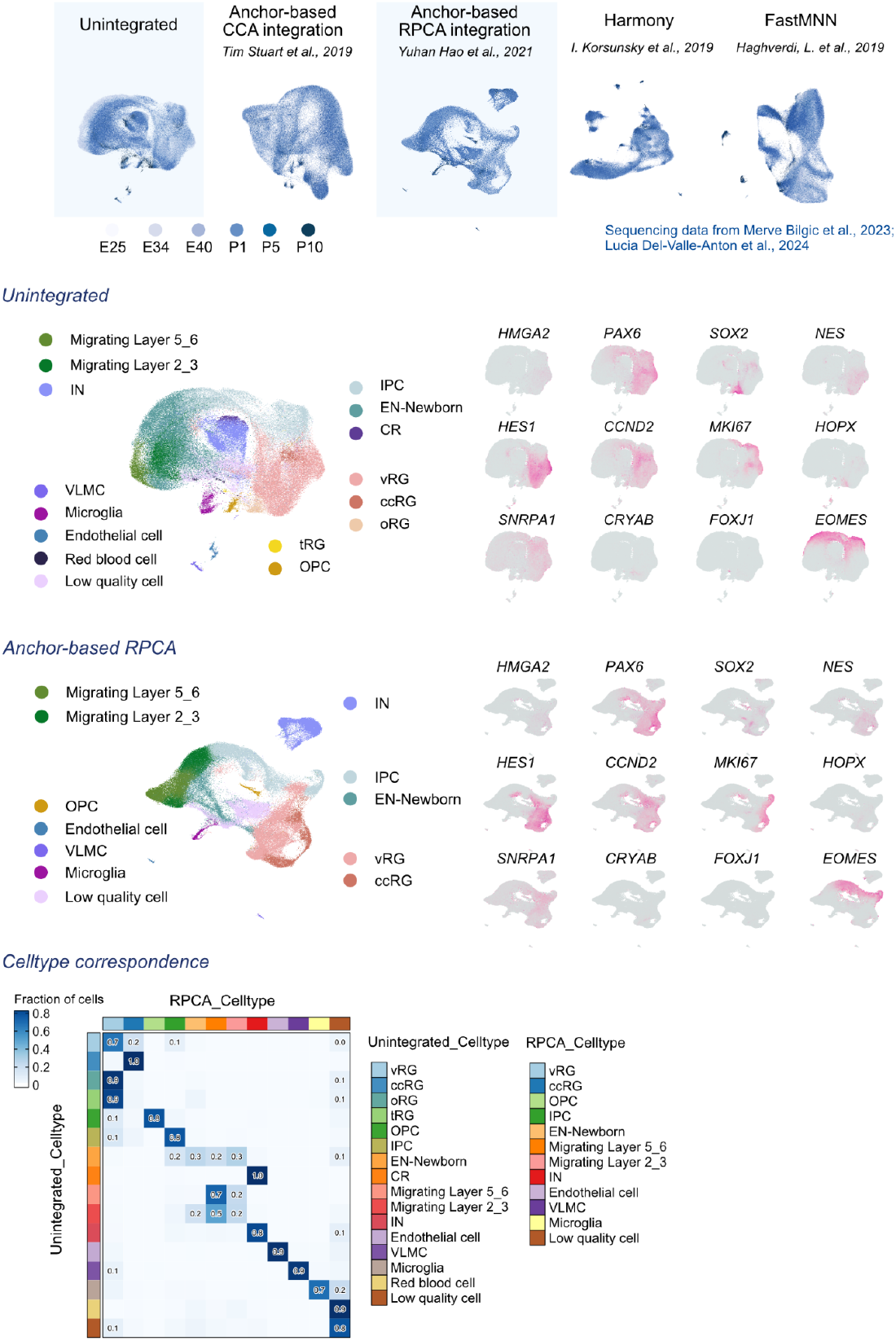
Cross-Validation of Cell Type Annotations in the Developing Ferret Cerebral Cortex. Unintegrated (SCT from Seurat) and RPCA methods were selected and assessed for marker expression fidelity. Correspondence of cell type assignments across integration methods was calculated and visualized along developmental stages. Based on consistent marker expression, the unintegrated method (default regression provided by Seurat) was chosen for cell type annotation in the ferret dataset.

**Figure S4:**
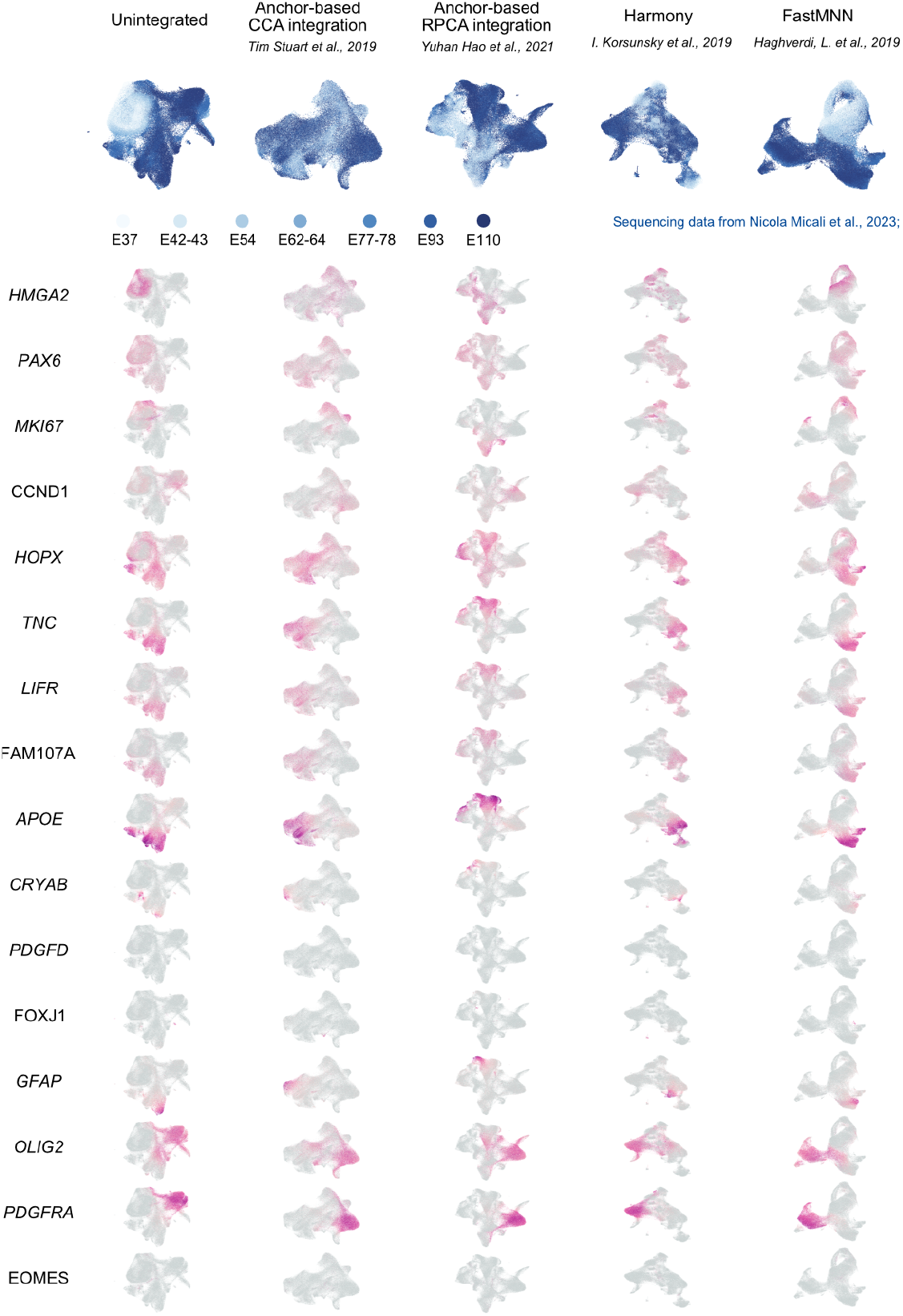
Categorization of Progenitor Cell Types in the Developing Macaque Cerebral Cortex. Five integration methods were applied to combine single-cell RNA-seq data from multiple developmental stages into a unified dataset, using canonical marker genes defined in this study. In the UMAP embedding space, cells are color-coded from light to dark blue to represent progression from early to late developmental stages. Expression patterns of canonical marker genes are visualized below the UMAP plots.

**Figure S5:**
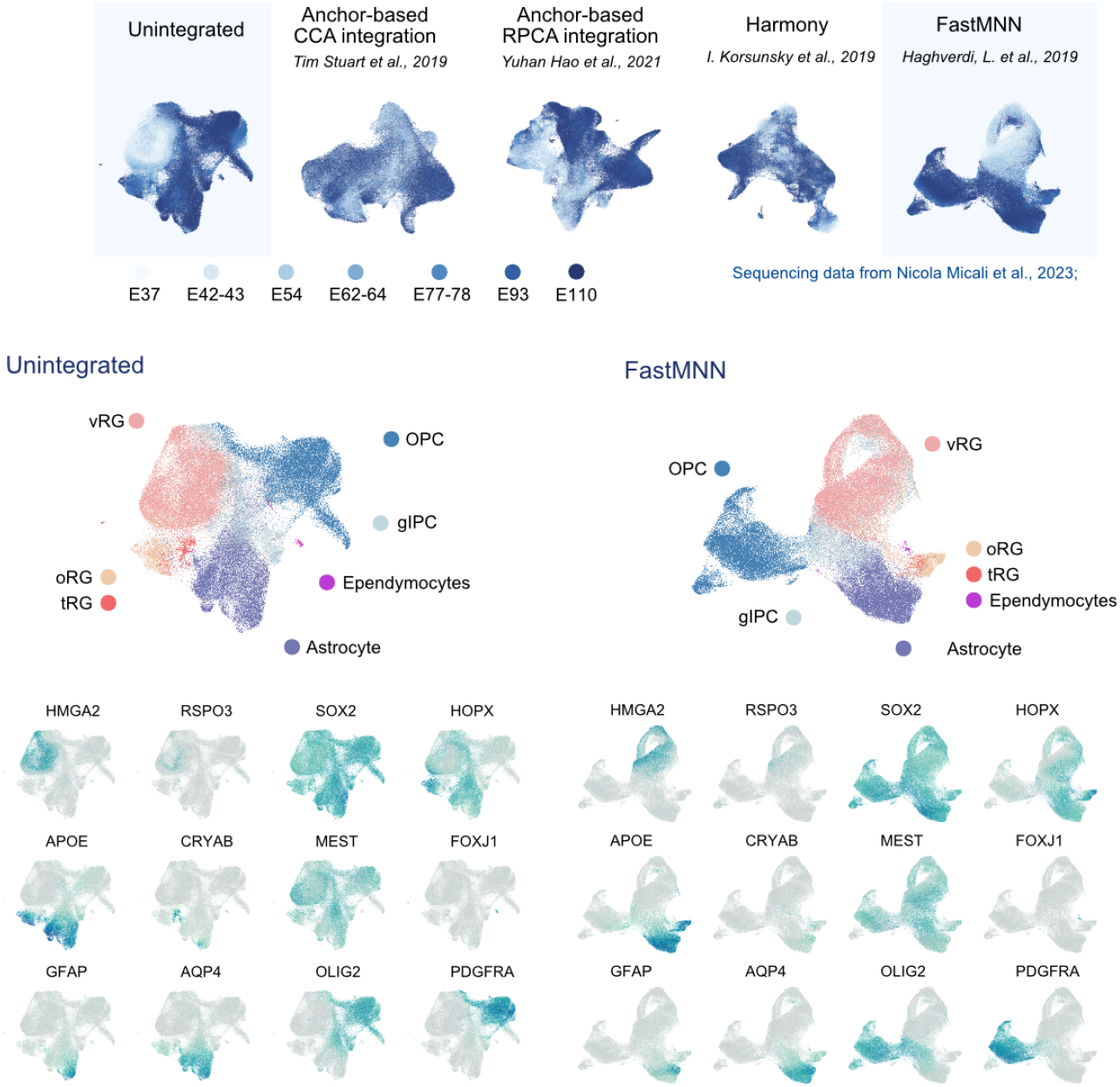
Reintegration of Progenitor Populations from Subsetted Macaque Datasets. To preserve accurate cell population distributions after subsetting progenitor cells (excluding differentiated neurons, oligodendrocytes, ependymocytes, and mature astrocytes based on marker expression and sampling stage), the dataset was re-regressed and integrated across developmental stages using five integration methods, consistent with prior analyses. Unintegrated (SCT from Seurat) and FastMNN methods were evaluated for marker expression fidelity. FastMNN was selected for determining progenitor cell types in the macaque dataset from the results section.

**Figure S8:**
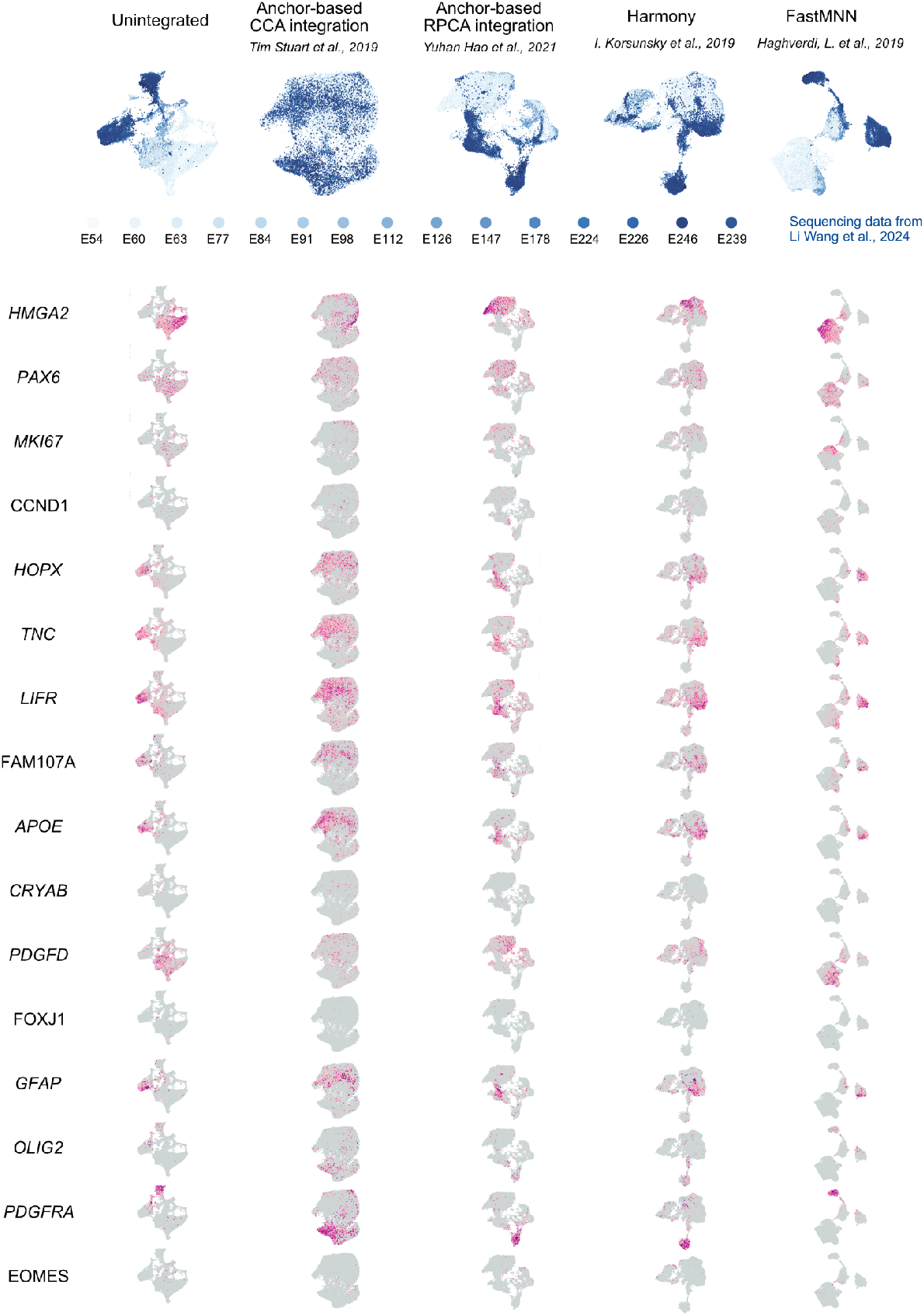
Categorization of Progenitor Cell Types in the Developing Human Cerebral Cortex. Five integration methods were applied to combine single-cell RNA-seq data from multiple developmental stages into a unified dataset, using canonical marker genes defined in this study. In the UMAP embedding space, cells are color-coded from light to dark blue to represent progression from early to late developmental stages. Expression patterns of canonical marker genes are visualized below the UMAP plots.

**Figure S9:**
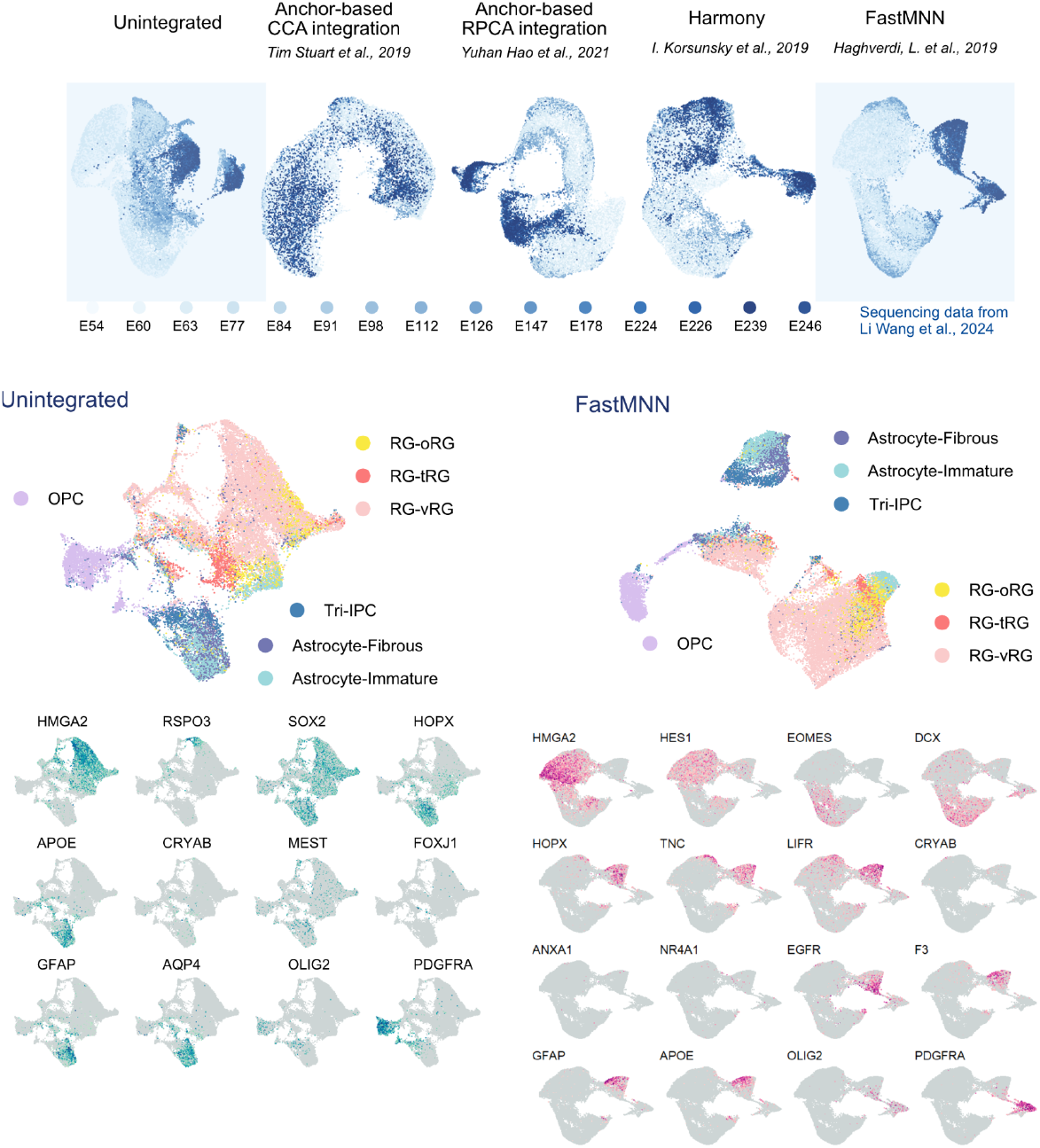
Reintegration of Progenitor Populations from Subsetted Human Datasets. To preserve accurate cell population distributions after subsetting progenitor cells (excluding differentiated neurons, oligodendrocytes, ependymocytes, and mature astrocytes based on marker expression and sampling stage), the dataset was re-regressed and integrated across developmental stages using five integration methods, consistent with prior analyses. Unintegrated (SCT from Seurat) and FastMNN methods were evaluated for marker expression fidelity. FastMNN was selected for determining progenitor cell types in the results section.

**Figure S10.**
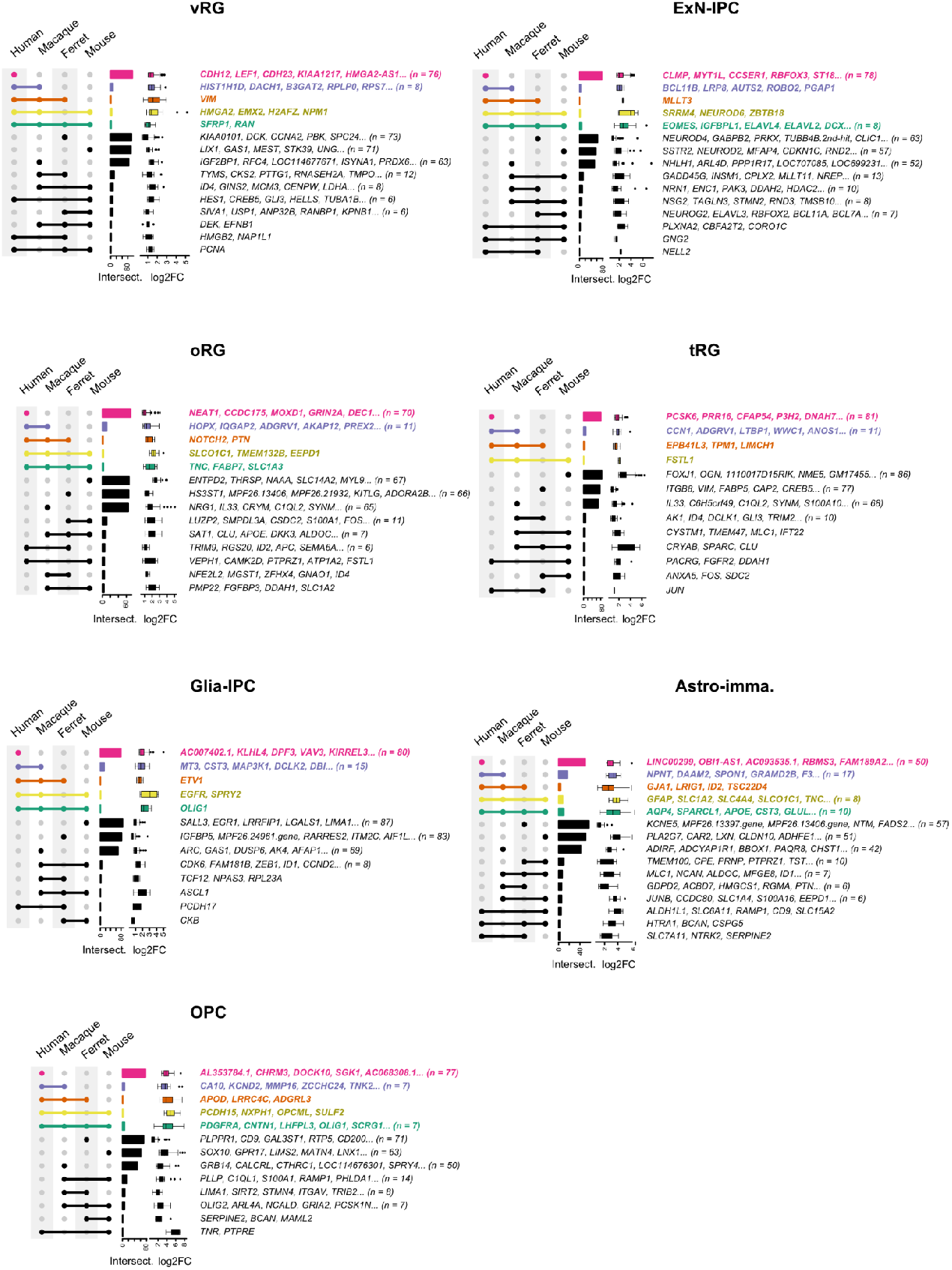
Marker Gene Conservation and Divergence Across Mammalian Species. UpSet plot illustrating the overlap of marker genes (top 100 per cell type, selected by differential expression analysis and ranked by log fold change) across four mammalian species. Color coding denotes specificity: pink (human-specific), purple (primate-specific), orange (gyrification-associated species), yellow (Euarchontoglires-specific), and green (conserved across all four species). Overlap patterns are calculated for each cell type documented in the study.

**Figure S11.**
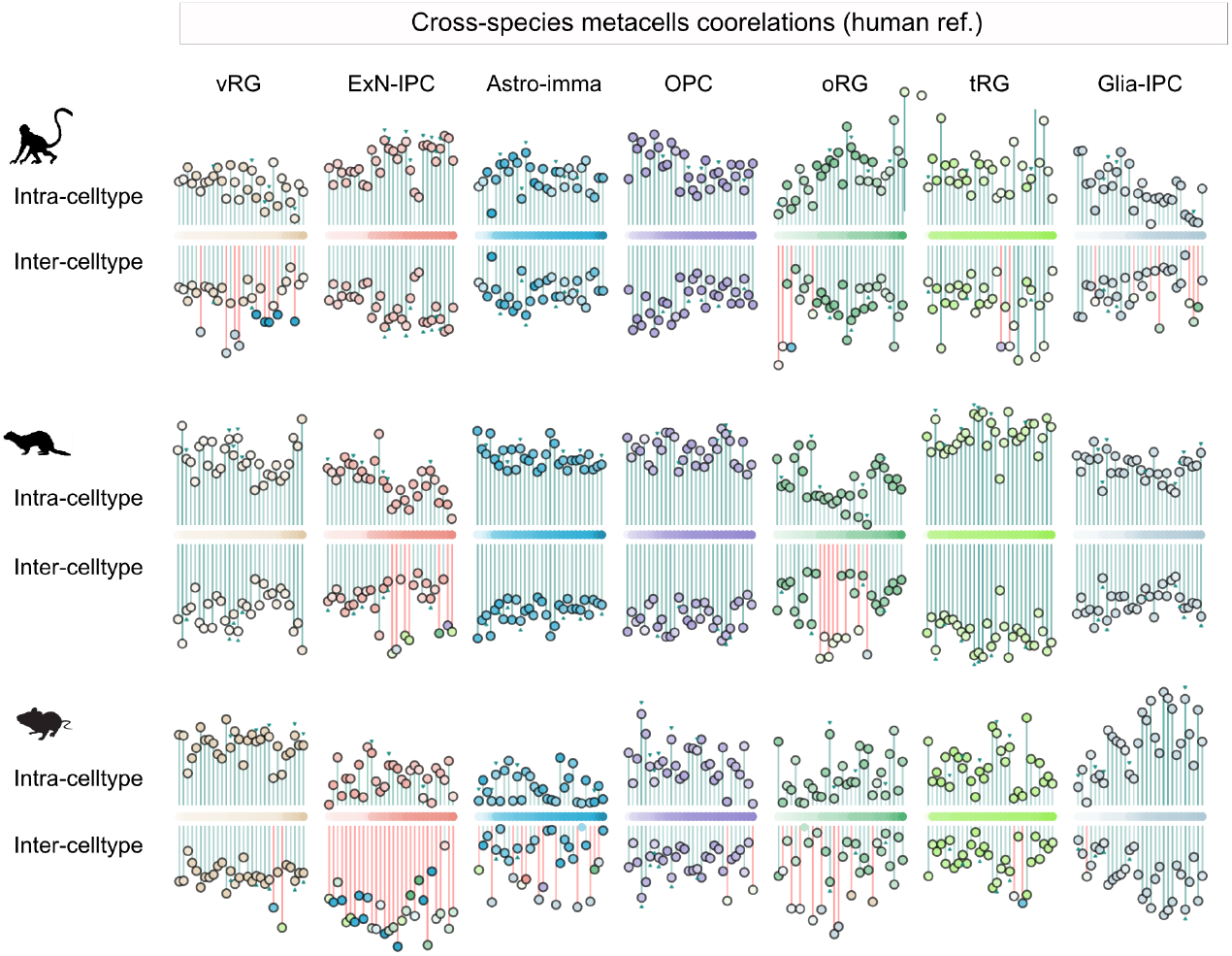
Cross-Species Correlations of Reciprocal Best Orthologous Metacells. Metacells are computed for each cell type, with cell clusters partitioned into a comparable number of metacells (cell assemblies with homogeneous gene expression, see methods). Metacells are colored based on the average developmental stage of their constituent cells, transitioning from light to dark shades. Each panel comprises two sections: (1) Intra-cell type: Identifies the local maximum correlation metacell pair within the same cell type between human and another species. (2) Inter-cell type: Identifies the global maximum correlation metacell pair across different cell types between human and other species. The distance from the panel center to each metacell represents the correlation value (ranging from 0, dissimilar, to 1, similar).

**Figure S12.**
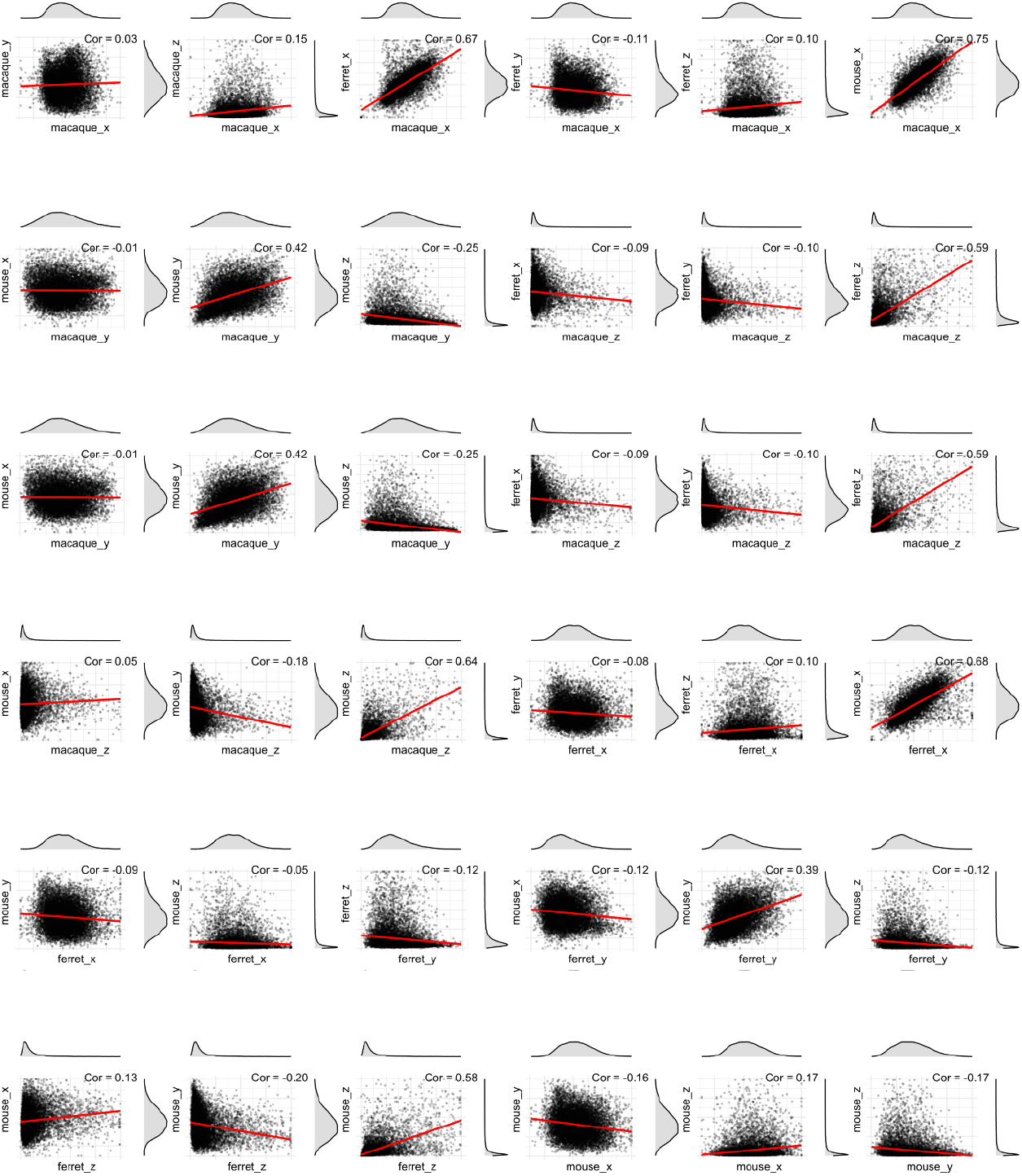
Pairwise comparisons of X, Y, and Z metrics (n = 36) with linear regression assessed correlation. Axis values from the three-dimensional gene expression divergence space were extracted and used to compute pairwise covariance correlations to assess relationships between axis metrics, both within and across species. Metric distributions are displayed on the right and top sides of the plot, corresponding to the y-axis and x-axis, respectively. A red line indicates the linear regression function, with the R-squared value annotated on the plot.

**Figure S13.**
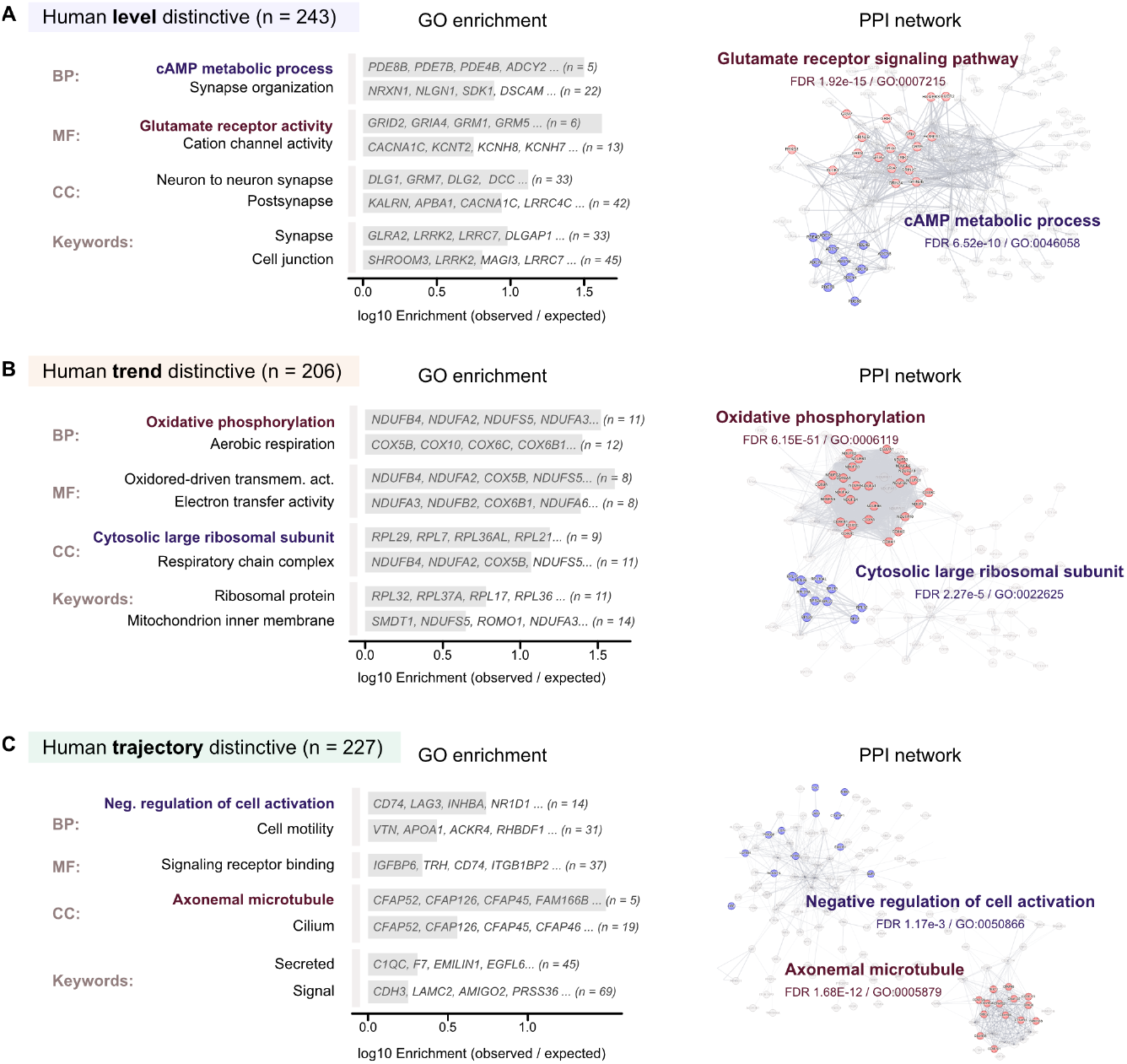
PGene Ontology Enrichment Analysis of HLD, HTD, and HDD Gene Sets. (A–C) Each panel presents a Gene Ontology (GO) enrichment plot for the HLD, HTD, and HDD gene sets, respectively. GO terms are categorized into Biological Process (BP), Molecular Function (MF), Cellular Component (CC), and UniProt Protein Keyword Annotations (Keywords). The x-axis represents the log10 enrichment score (observed/expected ratio). On the right, a protein-protein interaction (PPI) network derived from STRING analysis is shown. The top two enriched terms in each plot are highlighted in red and blue.

**Figure S14.**
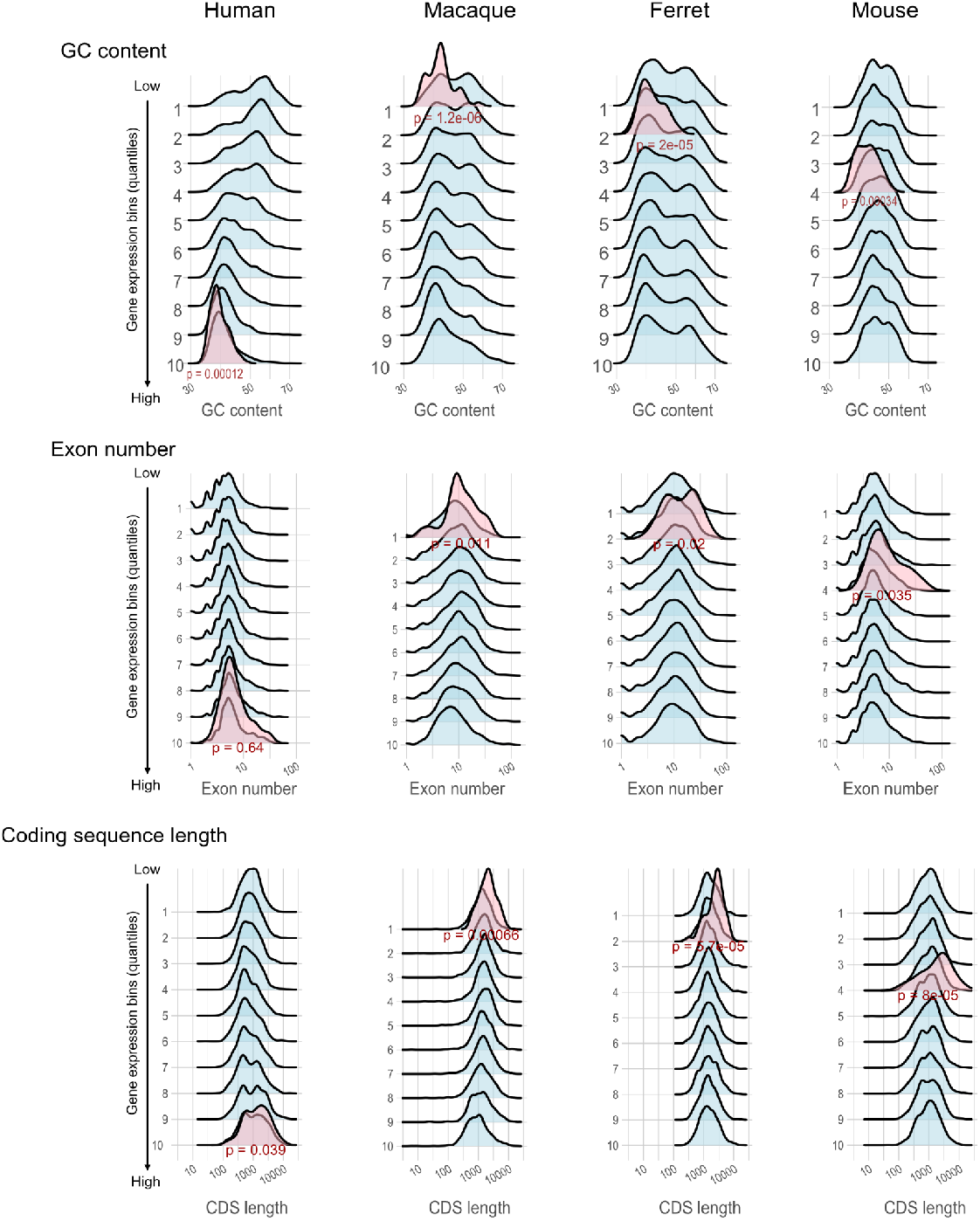
Distribution of GC contents, Exon number and Coding sequence length for homologous genes. All homologs (n=10683) genes were binned based on their expression magnitude. The distribution of GC contents, Exon number and Coding sequence length for HLD genes within each bin was visualized in pink to assess variations across expression levels. Statistical significance of differences in gene length distributions between HLD genes and a reference gene set (e.g., background or conserved genes) was evaluated using a corrected Kolmogorov–Smirnov test, with p-values reported to indicate the likelihood of observed differences arising by chance.

**Figure S15.**
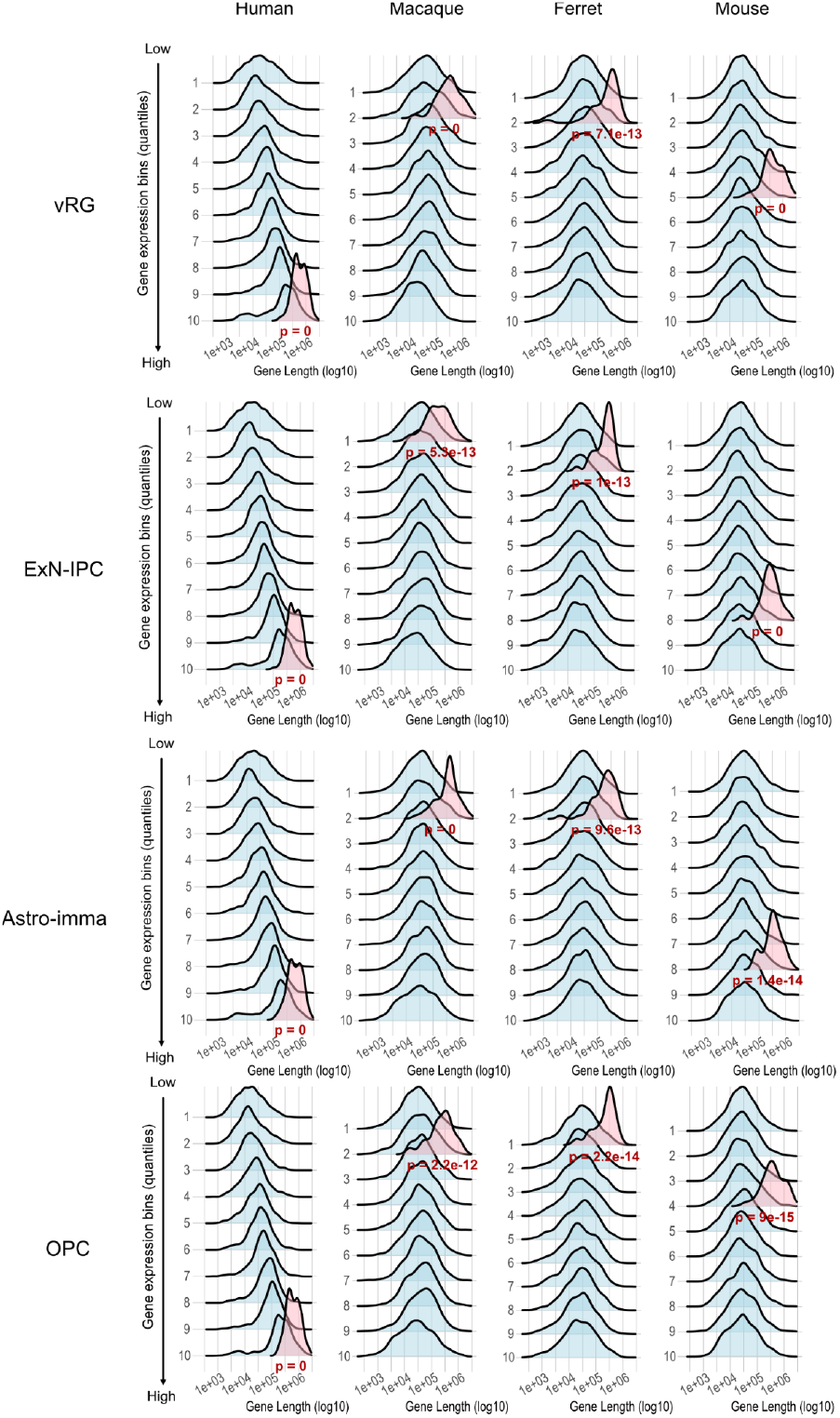
Cell-Type-Specific Genomic Span Analysis of HLD Genes Across Expression Levels. For each cell type, gene expression levels were quantified, and all annotated homologous genes (n=10683) were binned based on their expression magnitude. The distribution of genomic span lengths (including introns and exons) for HLD genes within each bin was visualized to assess variations across expression levels and cell types. Statistical significance of differences in gene length distributions between HLD genes and a reference gene set (e.g., background or conserved genes) was evaluated using a corrected Kolmogorov–Smirnov test, with p-values reported to indicate the likelihood of observed differences arising by chance.

**Figure S16.**
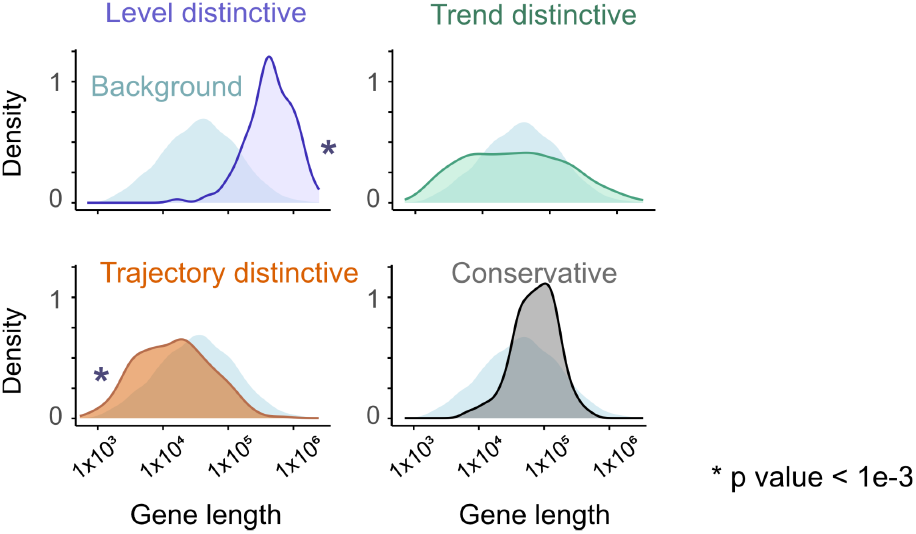
Gene lengths of HLD, HTD, and HDD classes. Three human distinctive gene sets are plots along with all homologs (n=10683). Statistical significance of differences in gene length distributions between gene set and a reference gene set (e.g., background or conserved genes) was evaluated using a corrected Kolmogorov–Smirnov test, with p-values reported to indicate the likelihood of observed differences arising by chance.

**Figure S17.**
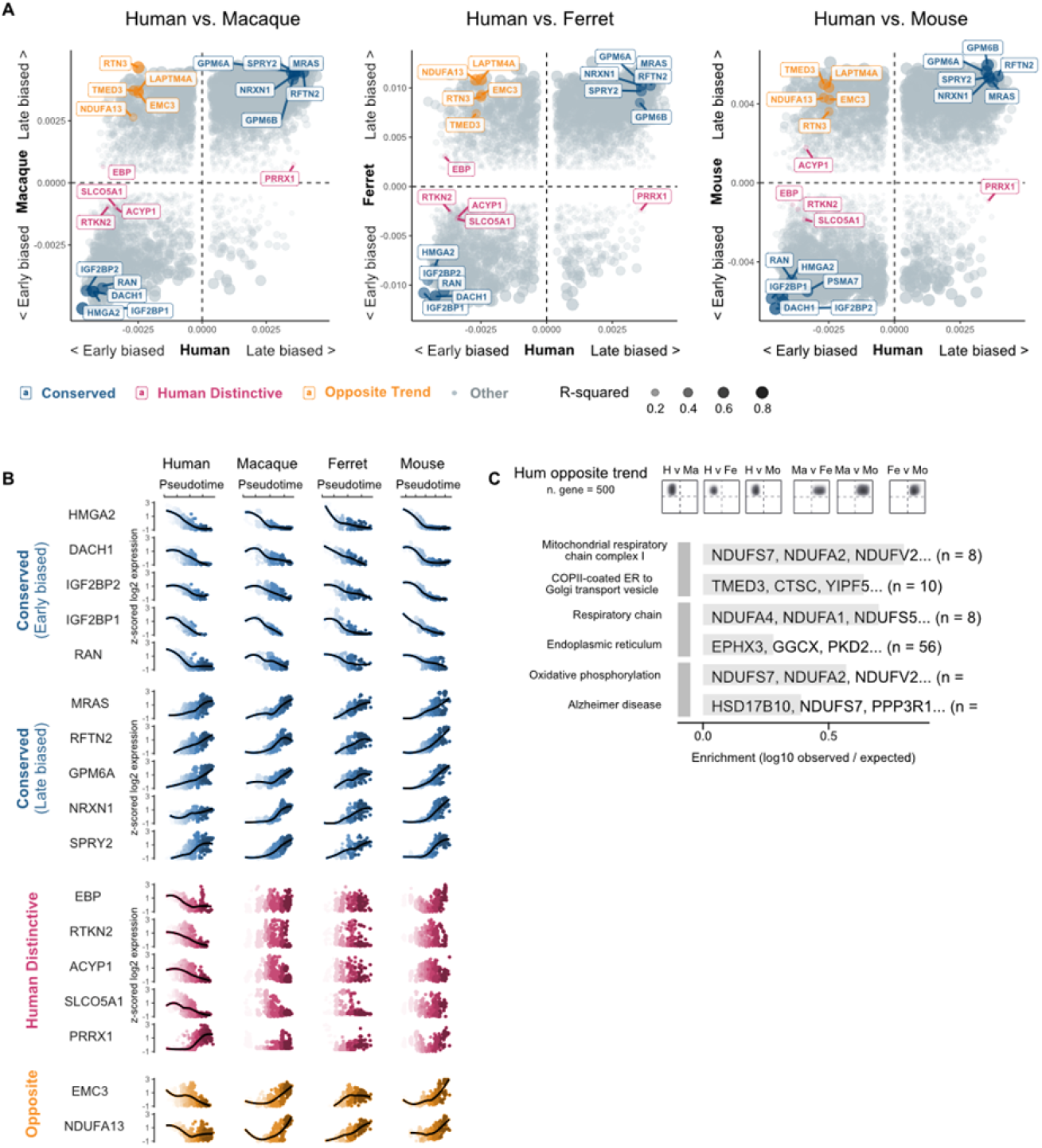
Gene Expression Trend Conservation and Divergence Across Species. **(A)** Gene sets categorized by linear regression coefficient patterns: conserved trends (consistent coefficient sign across all species, highlighted in blue), opposite trends (human coefficient sign opposes other species, highlighted in orange), and human-distinctive trends (temporal trend in human with near-zero coefficients in other species, highlighted in pink). Dot size represents the R-squared value. **(B)** Detailed visualization of identified genes from (A), plotted along the pseudotime axis for each species to illustrate expression dynamics. **(C)** Gene Ontology (GO) enrichment analysis of genes with opposite trends, as depicted in the accompanying density plot.

**Figure S18.**
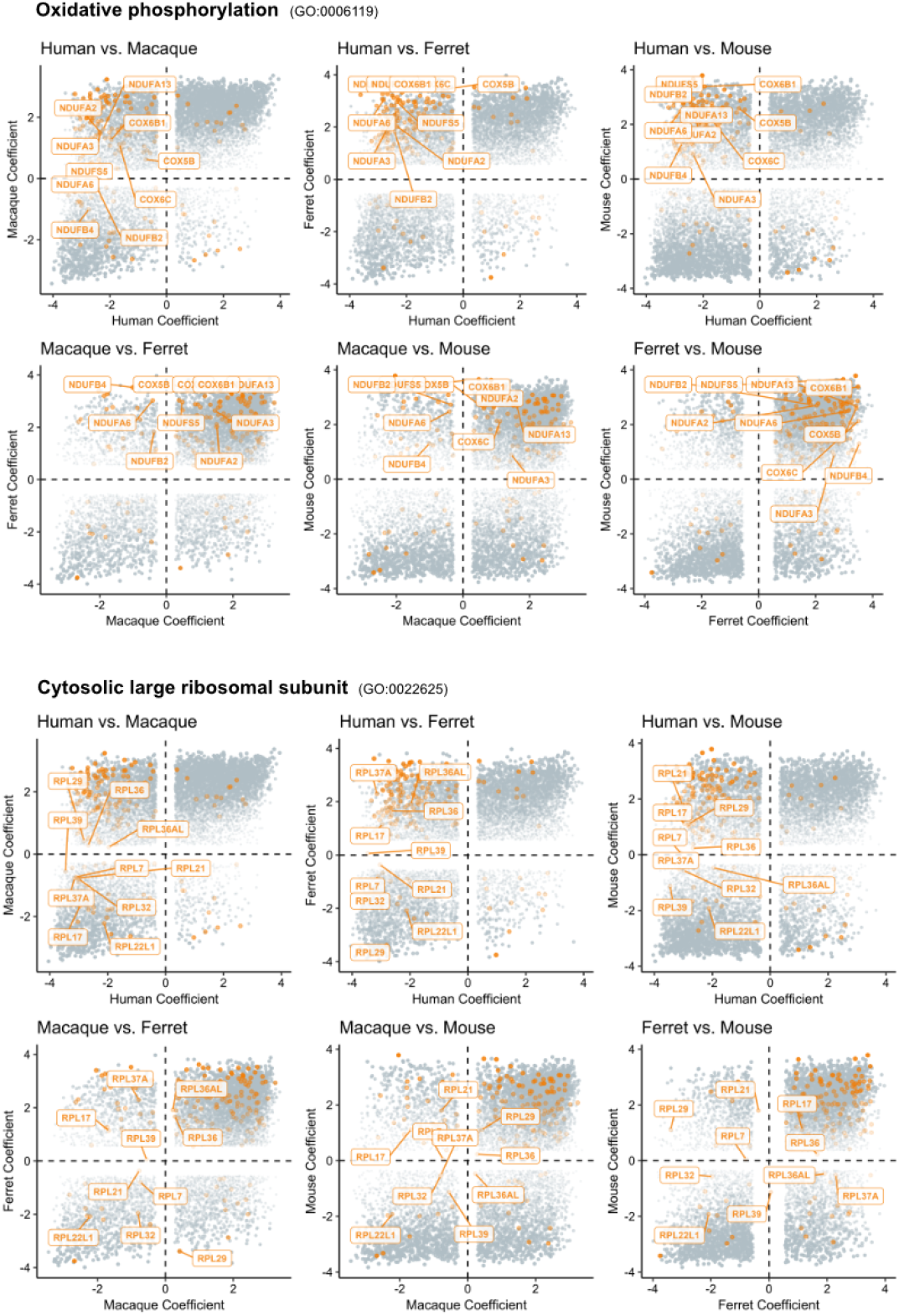
Cross-Species Coefficient Distributions of Genes in Highlighted GO Terms. Six pairwise comparisons of gene expression trend coefficients derived from linear regression across species, plotted with kernel-estimated gene density distributions. Genes associated with highlighted Gene Ontology (GO) terms (Oxidative Phosphorylation and Cytosolic Large Ribosomal Subunit) are labeled as yellow dots with gene names. Grey dots represent the background gene set (all genes).

**Figure S19.**
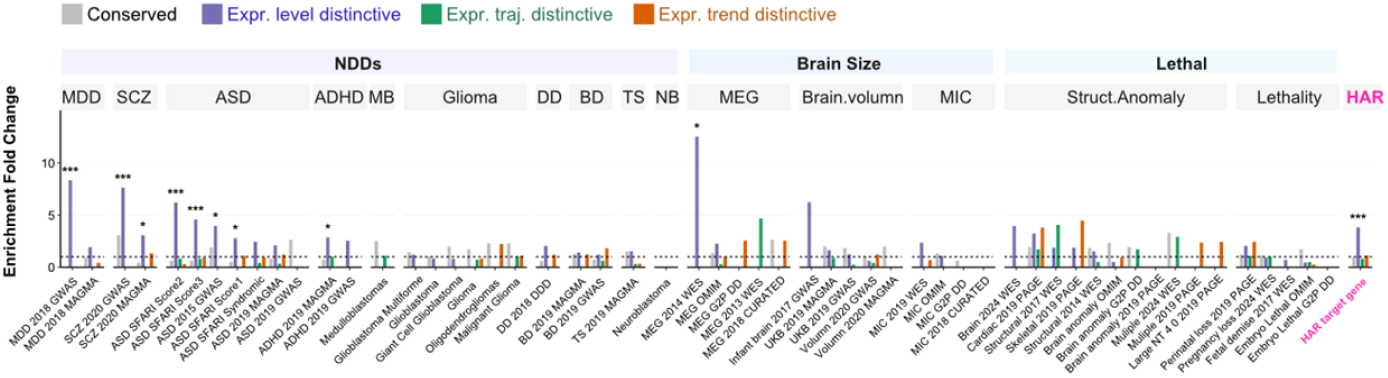
Disease enrichment analysis of human level-, trend-, and trajectory-distinctive gene sets. Enrichment is calculated as the ratio of observed to expected gene counts, adjusted for gene set size. Disease-associated genes were curated using the same methodology as in Figure 16. Among the three gene sets—Human Level Distinctive (HLD), Human Trend Distinctive (HTD), and Human Differentiation Trajectory (HDD)—the HLD set exhibits consistent enrichment across multiple neurodevelopmental disorders.

## Methods

### Trend Dissimilarity

To analyze how gene expression trends differ between species across neural developmental stages, we modeled the mean expression profile for each gene (*g*) using a Generalized Additive Model (GAM). This was done separately for human (denoted as species_1_) and a compared species (denoted as species_2_), which could be macaque, ferret, or mouse). GAMs are particularly useful here because they allow us to capture non-linear patterns in expression over time without assuming a specific functional form, providing a flexible yet robust fit to the data.

The GAM model for a given species (*k*) is expressed as

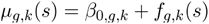

where:

(*µ*_*g,k*_ (*s*)) represents the expected expression level of gene (*g*) at a normalized developmental stage (*s*) (ranging from 0 to 1) for species (*k*), (*β*_0,*g,k*_) is the intercept term specific to gene (*g*) and species (*k*), (*f*_*g,k*_ (*s*)) is a smooth spline function of (*s*), constructed with up to 10 knots. This knot limit balances flexibility in capturing trends with the risk of overfitting noisy data.

Using these fitted models, we generated predicted expression values on a uniform grid of 100 points, defined as 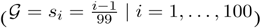. This grid ensures consistent comparison across species by sampling the developmental timeline evenly. The resulting prediction vectors are:

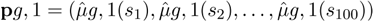

for human, and:

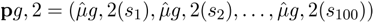

for the compared species. These vectors represent the smoothed expression trajectories over development.

To quantify the dissimilarity between these trajectories, we calculated the Euclidean distance (*d*_*g*_) between the two vectors:

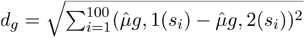

This distance measures the overall difference in expression trends across the developmental timeline. To make these distances comparable across genes and species pairs, we applied min-max normalization, scaling (*d*_*g*_) to a range of [0,1] for each species comparison:

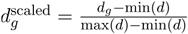

Here, min(*d*) and max(*d*) are the minimum and maximum *d*_*g*_ values observed across all genes within a specific species comparison, ensuring that the scaled metric reflects relative dissimilarity.

### Trajectory Bias Calculation Using Jensen-Shannon Divergence (JSD)

To evaluate how gene expression is distributed across different developmental trajectories, we used the Jensen-Shannon Divergence (JSD), a symmetric measure of difference between probability distributions. For each gene (*g*) and species (*k*), we computed the average expression across 11 distinct trajectories, forming a vector:

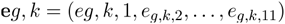

where (*e*_*g,k,j*_) is the mean expression of gene (*g*) in species (*k*) for trajectory (*j*). Since JSD requires probability distributions, we normalized these expression values into probabilities, adding a small constant (*ϵ* = 10^−6^) to avoid issues with zero values:

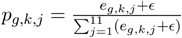

This ensures that 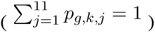, creating valid probability distributions (**p***g,k* = (*pg,k*,1, …, *pg,k*,11)) for each species.

The JSD between human ((*k* = 1)) and the compared species ((*k* = 2)) was then calculated as:

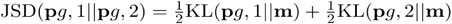

where 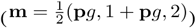 is the average distribution, and the Kullback-Leibler (KL) divergence is defined as:

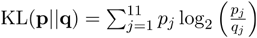

JSD provides a bounded measure in [0,1], where 0 indicates identical distributions and 1 indicates maximal divergence. This value directly serves as the trajectory bias score, reflecting differences in how gene expression is allocated across developmental paths between species.

### Log2 Fold Change (Log2FC) Calculation

To compare overall expression levels between species, we combined single-cell RNA sequencing data from humans and each compared species into merged Seurat objects. We applied SCTransform (SCT) to normalize the data and stabilize variance across genes, accounting for technical variability. Differential expression analysis was performed using a logistic regression (LR) test, with cell type included as a covariate to control for compositional differences.

The log2 fold change for gene (*g*) was calculated as:

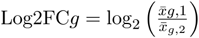

where 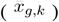 is the adjusted average expression of gene (*g*) in species (*k*) after SCT normalization. This metric quantifies the relative difference in expression magnitude, with positive values indicating higher expression in human and negative values indicating higher expression in the compared species.

### Identification of Human-Divergent and Human-Distinctive Genes

We integrated the three metrics—scaled Log2FC ((*x*_*g*_)), trend dissimilarity ((*y*_*g*_)), and trajectory bias ((*z*_*g*_)) —into a 3D space to classify genes. A gene (*g*) was classified as “human-divergent” for a given species comparison if it met one of these criteria:

The Euclidean distance from the origin in this 3D space exceeded the 95th percentile across all genes:

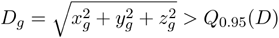

Or any single parameter exceeded its 98th percentile:

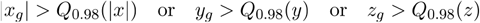

These thresholds identify genes with extreme differences in expression level, trend, or trajectory distribution. Genes were further classified as “human-distinctive” if they were identified as divergent in all three species comparisons (macaque, ferret, mouse), indicating consistent divergence from non-human species.

### Parameter Scaling

To ensure comparability across the three metrics, we applied the following scaling procedures:

#### Trend Dissimilarity

Scaled to [0,1] within each species comparison:

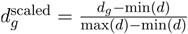

This normalization preserves relative differences while standardizing the range.

#### Trajectory Bias (JSD)

Used directly without additional scaling, as JSD is naturally bounded in [0,1], providing an inherent standardized metric.

#### Log2FC

Log2FC values across all species comparisons were first winsorized to the 0.1% and 99.9% quantiles to reduce the impact of extreme outliers. Then they were divided by the maximum absolute winsorized value (*M*) to get the scaled Log2FC range from -1 to 1:

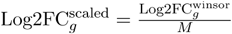

This scaling ensures that the distribution’s zero point (no change) is preserved, and extreme values are moderated, facilitating cross-species comparisons.

### Disease-derived vector in divergent space

To evaluate whether a disease gene set (e.g., ASD-associated genes) exhibits distinctive gene expression differences between humans and another animal (macaque, ferret, or mouse), we calculate the deviation vector derived from each disease in cross-species gene expression divergent space.For a given comparison, let G be the set of all genes, with each gene *i*∈**G** as a point P*i* = (*x*_*i*_,*y*_*i*_,*z*_*i*_).

The species vector is computed as:

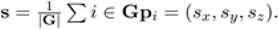

For a disease gene set **D**⊆**G**, the disease vector is:

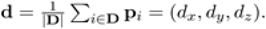

The deviation vector, capturing the disease set’s departure from the species average, is:

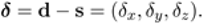

To test if ***δ*** is significantly aligned with an axis *k* (where *k* = *x,y,z*), we compute the angle *θ*_*δ,k*_ between ***δ*** and the unit vector *e*_*k*_ (e.g., ): *ex* = (1,0,0)

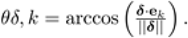

Significance is assessed via a permutation test: 10,000 random gene sets **R**_*j*_ of size |**D**| are sampled from **G**, with deviation vectors *δr*_*j*_ = r*j* − s, where 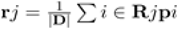. The angle for each is 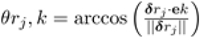. The p-value is:

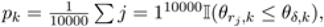

where I is the indicator function. A small *p*_*k*_ < 0.05 indicates the disease gene set’s deviation is significantly aligned with axis *k*, suggesting distinctiveness in level (x), trend (y), or trajectory (z). This approach was applied across human vs. macaque, ferret, and mouse comparisons for various disease gene sets

To visualize the deviation of disease-associated gene sets from the species average in gene expression divergent space, we plotted the components of the deviation vector (x, y, z) as horizontal bars (Figure 6 C-H). Each bar extends from zero to the observed deviation value, with the zero point centered in the plot. Ticks indicate the 95% range of permutation-based background distributions (n = 10,000) and the zero point. Observed values and Bonferroni-adjusted p-values (with significance stars) are labeled near the bar, positioned to remain within the plot boundaries. The plots are vertically aligned with consistent scaling to facilitate comparison across components.

### Evaluate axes contribution to total variance by extreme gradient boosting

To evaluate the contributions of divergence metrics (*x, y*, or *z*) ---representing differences in expression level, trend, and trajectory, respectively---to the variance in gene expression between species, we employed the Extreme Gradient Boosting (from xgboost r package) algorithm. XGBoost was selected due to its superior capability to model complex, non-linear relationships and interactions among variables, which is critical for capturing the subtle and interdependent effects of these metrics on gene expression variance.

For each pairwise species comparison (e.g., human vs. macaque, human vs. ferret, human vs. mouse), we first calculated the between-species variance per gene. This was defined as the squared difference in mean expression levels between the two species.

We trained an XGBoost regression model to predict between-species variance using the divergence metrics (*x, y*, or *z* ) as input features. The model’s hyperparameters were carefully tuned to optimize predictive accuracy while preventing overfitting and ensuring generalizability across diverse species comparisons.

The key parameters were set as follows:

#### Objective function

Regression with squared error loss, minimizing the difference between predicted and actual variance values.

#### Maximum tree depth

*6*, restricting the depth of each decision tree to limit model complexity and reduce the risk of overfitting to noise in the data.

#### Learning rate (η)

0.3, determining the step size at which the model updates its predictions with each boosting iteration, balancing convergence speed and stability.

#### Number of boosting rounds

100, specifying the total number of trees in the ensemble, sufficient to capture feature effects without excessive computation.

#### Subsample ratio

0.8, meaning that 80% of the training data were randomly sampled to build each tree, enhancing robustness by reducing sensitivity to individual data points.

#### Column subsample ratio

0.8, indicating that 80% of the features (*x, y*, or *z*) were randomly selected for each tree, introducing additional randomness to prevent overfitting and improve generalization.

The training process involved a 5-fold cross-validation scheme to assess model performance and ensure stability across different subsets of the data. This step confirmed that the chosen hyperparameters were effective in avoiding overfitting while maintaining predictive power.

The total variance explained by the full model (including all three metrics) was quantified using the coefficient of determination 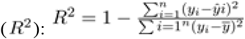 where:

(*y*_*i*_) is the actual between-species variance for gene ( i ),

(*ŷ*_*i*_) is the variance predicted by the XGBoost model,

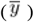 is the mean variance across all genes,

(*n*) is the total number of genes analyzed.

An (*R*^2^) value close to 1 indicates that the metrics ( x ), ( y ), and ( z ) collectively explain most of the observed variance, while a value near 0 suggests limited explanatory power.

To isolate the unique contribution of each metric (*x, y*, or *z*) to the explained variance, we adopted a subtraction-based approach. For each metric, we trained a reduced XGBoost model excluding that metric (e.g., using only ( y ) and ( z ) when excluding ( x )) and recalculated the ( R^2 ) for the reduced model. The partial contribution of the excluded metric was then computed as:

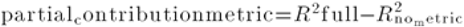

where:

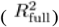 is the ( R^2 ) from the model with all three metrics,

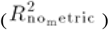 is the ( R^2 ) from the reduced model excluding the metric of interest.

This difference reflects the additional variance explained by the excluded metric beyond what the remaining metrics can account for. The process was repeated for each metric (*x, y*, or *z*), providing a clear breakdown of their individual impacts.

### Histology and immunostaining

Mouse embryos were collected after electroporation and perfused with ice-cold 4% paraformaldehyde (PFA) in PBS. Dissected brains were soaked in 4% PFA overnight at 4°C and sectioned into 100 μm slices using a vibratome (Leica, Cat# VT1000S). Slices were washed with PBST (PBS with 0.1% Triton X-100) three times and incubated in blocking solution (PBS with 0.3% Triton X-100 and 3% donkey serum) for 30 minutes. Brain slices were then incubated overnight at 4°C with primary antibodies (Table 2). After three PBST washes, slices were incubated with secondary antibodies (Table 3) for 2 hours at room temperature. After additional PBST washes, slices were mounted on slide glasses using DAKO glycerol mounting medium (Cat# C0563). Imaging was performed with a confocal microscope (Olympus FV3000), and images were processed using Fiji/ImageJ software.

### Quantification and statistical analysis

Statistical analyses were performed using GraphPad Prism 10 (GraphPad Software, Inc., USA) and Microsoft Excel (Microsoft Inc., USA). Experiments were repeated at least three times, and results are expressed as mean ± SEM. Statistical significance of disease deviation vectors was assessed by comparing observed values to permutation-based background distributions (n=10,000). Bonferroni-adjusted p-values are annotated with significance stars adjacent to the corresponding bars.

### Supplementary tables

Supplementary Table 1-Human distinctive genes

Supplementary Table 2-Coordinate of homologs gene in divergent space Supplementary

Table 3-Curated disease associated genes

